# Phosphatidylethanolamine made in the inner mitochondrial membrane is essential for yeast cytochrome *bc*_1_ complex function

**DOI:** 10.1101/269233

**Authors:** Elizabeth Calzada, J. Michael McCaffery, Steven M. Claypool

**Affiliations:** Department of Physiology, Johns Hopkins University School of Medicine, Baltimore, MD, USA; Integrated Imaging Center, Department of Biology, Johns Hopkins University, Baltimore, MD

## Abstract

Of the four separate PE biosynthetic pathways in eukaryotes, one occurs in the mitochondrial inner membrane (IM) and is executed by phosphatidylserine decarboxylase (Psd1p). Deletion of Psd1, which is lethal in mice, compromises mitochondrial function. We hypothesize that this reflects inefficient import of non-mitochondrial PE into the IM. To test this, we re-wired PE metabolism in yeast by re-directing Psd1p to the outer mitochondrial membrane or the endomembrane system. Our biochemical and functional analyses identified the IMS as the greatest barrier for PE import and demonstrated that PE synthesis in the IM is critical for cytochrome *bc*_1_ complex (III) function. Importantly, mutations predicted to disrupt a conserved PE-binding site in the complex III subunit, Qcr7p, impaired complex III activity similar to *PSD1* deletion. Collectively, these data demonstrate that PE made in the IM by Psd1p is critical to support the intrinsic functionality of complex III and establish one likely mechanism.

## INTRODUCTION

The evolutionary design of living cells has revealed redundancies in various metabolic pathways to promote survival. Importantly however, the sequestration of enzymes and their substrates into different membrane compartments allows for the enrichment and regulation of metabolite synthesis in regions of the cell where they are essential. Such is the case for the biosynthesis of the abundant membrane phospholipid, phosphatidylethanolamine (PE). In mammalian cells PE is synthesized by four separate pathways, three of which localize to the endoplasmic reticulum (ER) ^1^. A final pathway is dependent on phosphatidylserine decarboxylase (Psd1p) which is spatially isolated within the mitochondrial inner membrane (IM) ^2-5^. The predominant pathways for PE production include the Kennedy pathway, which synthesizes PE through the stepwise conjugation of CDP-ethanolamine to diacylglycerol (DAG), and the Psd pathway which utilizes phosphatidylserine (PS) as a substrate to generate PE ^1^. In multicellular organisms, preference for either the Kennedy pathway or the Psd pathway varies between tissues and cell types ^6^. Notably, deletion of either pathway is lethal during murine embryogenesis, highlighting the importance of PE generation in both the ER and mitochondrial compartments for development ^7, 8^.

Conservation of the Psd pathway from bacteria to humans likely reflects the endosymbiotic origin of mitochondria which in turn suggests that mitochondrial PS and PE metabolism has been preserved to optimize mitochondrial performance ^6^. Indeed, deletion of phosphatidylserine decarboxylase (*Pisd* in mouse and humans and *PSD1* in yeast) in eukaryotic cells decreases cellular growth, impairs oxidative phosphorylation, produces aberrant mitochondrial morphology, and diminishes PE levels in cells and mitochondria ^7, 9-12^. The Psd pathway is the predominant pathway for PE production in *Saccharomyces cerevisiae* and produces up to 70% of PE in the cell ^13^. In contrast to mammals, yeast additionally contain Psd2p which localizes to either Golgi or endosomal compartments ^5, 14^. Deletion of *PSD2* alone does not recapitulate any of the mitochondrial defects associated with loss of *PSD1*, further emphasizing the importance of the mitochondrial PE biosynthetic pathway ^5^. The combined absence of *PSD1* and *PSD2* produces a strain that is auxotrophic for exogenous ethanolamine supplementation which is used to generate PE through the Kennedy pathway. As the severity of PE depletion can be genetically and metabolically adjusted in yeast, this organism has served as an invaluable model for studying the importance of the mitochondrial Psd pathway for cellular respiration.

Characterization of the Psd pathway over more than 50 years of research has revealed detailed mechanistic insight into the biogenesis of the mature Psd1p enzyme ^15^. In yeast, *PSD1* is a nuclear encoded gene whose transcript is translated on cytosolic ribosomes which produce a full-length zymogen that is catalytically inactive. Upon its mitochondrial import, two matrix metalloproteases sequentially process the N-terminus of Psd1p which then undergoes a third processing event that is executed internally by the enzyme itself at the conserved C-terminal LGST motif ^2, 3^. This unique autocatalytic processing event severs Psd1p into a large membrane-anchored β subunit and a small *α* subunit and is performed by a self-contained catalytic triad typical of serine proteases ^16, 17^. Self-processing endows the small *α* subunit with an N-terminal pyruvoyl prosthetic group that is crucial for its decarboxylase activity. Post-cleavage, the two Psd1p subunits remain non-covalently associated within the inner membrane where they decarboxylate PS to PE ^17^.

The substrate of Psd1p, PS, is synthesized on the mitochondrial-associated membrane (MAM) of the ER by phosphatidylserine synthase (Cho1p) ^18^. Thus, the amphipathic PS must be imported from the ER and traverse through two aqueous compartments, the cytosol and the mitochondrial intermembrane space (IMS), to reach Psd1p at the IM ^19^. While the redundant roles played by ER-mitochondria membrane tethers in PS mitochondrial import continue to be elucidated ^20-24^, recent evidence in yeast indicates that Mic60p, a subunit of the mitochondrial contact site and cristae organizing system (MICOS), works in conjunction with the soluble lipid carrier, Ups2p, to expose PS for Psd1p decarboxylation in the IMS ^25, 26^. Whether or not a parallel pathway exists for PE import into the IM remains unclear. The lethal consequence of *pisd* deletion in mice and the failure of supplemental ethanolamine to rescue the respiratory defects of *psd1*Δ yeast were taken as evidence that PE made outside of the mitochondrion is unable to compensate for the absence of Psd1p ^1, 7, 11, 27^. However, it was recently reported that PE made by the Kennedy pathway can in fact restore the impaired oxidative phosphorylation of *psd1*Δ yeast suggesting that extra-mitochondrial PE can compensate for the absence of the Psd pathway ^28^.

Here, we sought to determine if either the cytosol and/or the IMS is a barrier that prevents PE made outside of the IM from rescuing the impaired oxidative phosphorylation that occurs in *S. cerevisiae* lacking Psd1p. Previously, we generated a chimeric Psd1p protein, ER-Psd1p, which is targeted to the endosomal compartment and is catalytically active ^29^. In the current study, we further characterized the cell biology of ER-Psd1 yeast, together with an OM-targeted chimeric Psd1p (OM-Psd1p), to test if the cytosol, IMS, or both, are barriers that prevent non-mitochondrially produced PE from functionally rescuing the absence of PE made in the IM. Alongside strains expressing these re-directed Psd1p constructs, we compared the mitochondrial function of *psd1*Δ*psd2*Δ yeast grown in the non-fermentable carbon source, lactate, with or without exogenous ethanolamine supplementation, to evaluate the ability of the ER-localized Kennedy pathway to support mitochondrial function. In contrast to cytochrome *c* oxidase (complex IV), cytochrome *bc*_1_ respiratory complex (complex III) activity was impaired in the absence of IM-synthesized PE even after supplementation with ethanolamine. Intriguingly, the positive effect of ethanolamine on complex IV activity reflected an increase in mitochondrial levels of cardiolipin (CL), and not PE. Notably, the absence of Cho1p disrupted complex III and IV activities without altering Psd1p steady state levels. These results highlight a crucial role of PE for complex III function and suggest that the IMS is the greatest obstacle preventing extra-mitochondrially produced PE from functionally substituting for IM-fabricated PE. Structures of yeast and human complex III have each revealed a bound PE in close proximity to the complex III subunit, Qcr7p ^30, 31^. Strikingly, mutations predicted to disrupt PE-binding by Qcr7p impaired complex III activity to a similar level as in the absence of Psd1p. Altogether, we demonstrate that PE made in the IM by Psd1p is critical to support full complex III function and provide the first molecular evidence of the functional importance of a PE-binding site in complex III.

## RESULTS

### The impaired respiratory growth of psd1Δ yeast is not rescued by PE synthesis through the Kennedy pathway

The ability of supplemental ethanolamine to rescue the respiratory growth defect of *psd1*Δ yeast has been reported by some groups ^28^ but not others ^1, 11, 27^. This is an important controversy to resolve. If extra-mitochondrially produced PE can replace PE normally made in the IM, then this would imply that the mitochondrial Psd1 pathway *per se*, is not necessary for mitochondrial function. By extension, it would suggest that mechanisms to move extra-mitochondrially produced PE into mitochondria are robust enough to compensate for the absence of Psd1p. Therefore, we tested the growth of wildtype (WT), *psd1*Δ, *psd2*Δ, and *psd1*Δ*psd2*Δ in synthetic complete ethanol-glycerol (SCEG) medium with or without 2mM ethanolamine supplementation (Fig 1A). Consistent with previous findings ^1, 11, 27^, we found that exogenous ethanolamine supplementation restored respiratory growth of *psd1*Δ*psd2*Δ yeast to *psd1*Δ levels but failed to fully restore the respiratory defect associated with *psd1*Δ yeast. Importantly, this basic result was confirmed in *psd1*Δ yeast made in three additional yeast strain backgrounds (Fig 1B, note that the supplemental ethanolamine concentration was increased to 10mM), although subtle differences between the strains were noted. Overall, these findings indicate that the Kennedy pathway cannot fully compensate for the absence of Psd1p function, potentially due to inefficient trafficking of PE from the ER to mitochondrial OM and/or from the OM to the IM.

**Fig 1.**
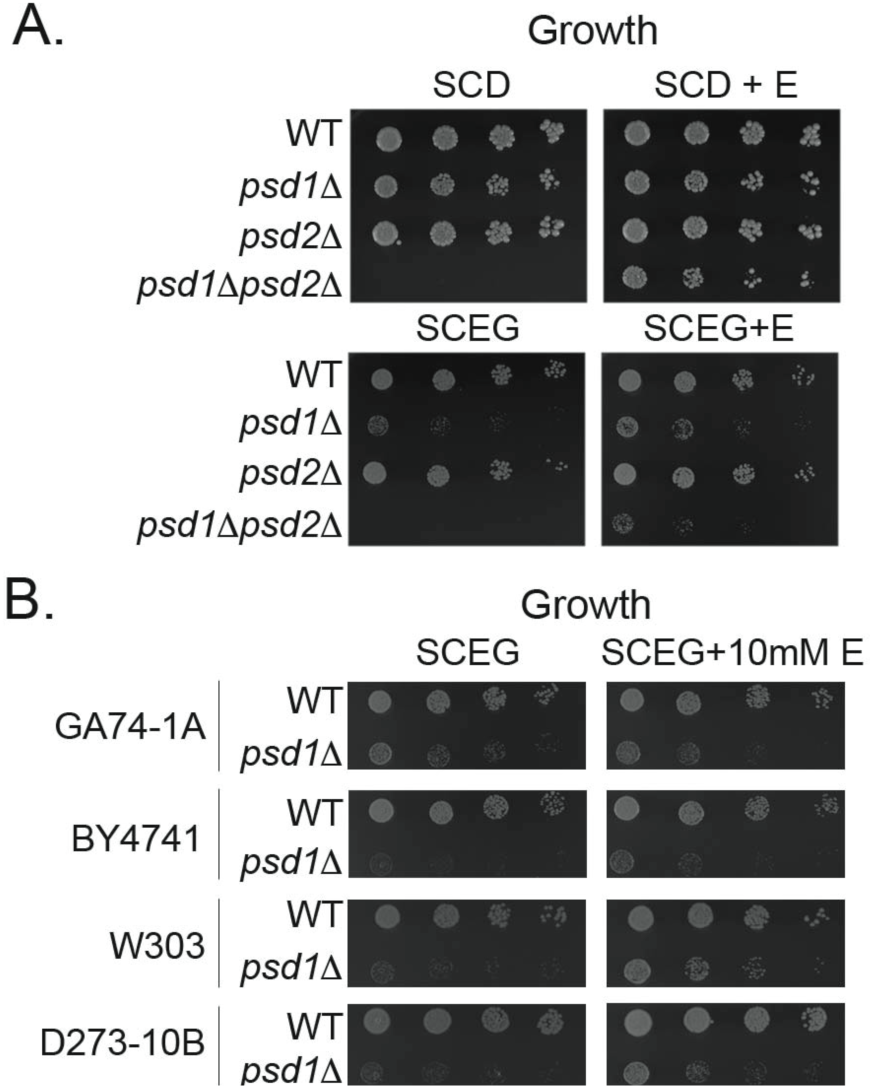
Ethanolamine supplementation fails to rescue the growth defect of *psd1*Δ on respiratory medium in 4 separate yeast strains. (A, B) The indicated strains were pre-cultured at 30°C in YPD and spotted onto (A) synthetic complete dextrose (SCD) or ethanol-glycerol (SCEG) medium +/- 2mM ethanolamine (+E) or (B) SCEG +/- 10mM ethanolamine and incubated at 30°C for 3 days.

### Validation of constructs that re-direct Psd1p to OM or ER membranes

To interrogate whether the cytosol and/or the IMS is a barrier that prevents extra-mitochondrially produced PE from replacing PE normally made in the IM, we generated chimeric Psd1p constructs that are localized to either the ER or OM to redirect PS and PE metabolism. As depicted in Fig 2A, these two constructs, and the WT IM-localized Psd1p control (referred to as IM-Psd1p to distinguish it from strains expressing endogenous Psd1p), contain a C-terminal 3XFLAG tag to track autocatalytic function of these chimeras by immunodetecting the released Psd1p α subunit. To re-direct Psd1p to the mitochondrial OM, the mitochondrial targeting sequence and transmembrane domain of Psd1p were replaced by the equivalent domains of the single-pass OM resident protein, Tom20p. ER-Psd1p, which is directed to the secretory pathway, was generated by replacing the mitochondrial targeting sequence of Psd1p with the NH_2_-terminal signal sequence of carboxypeptidase Y (CPY) ^29, 32^. The ER-Psd1p construct additionally contains an *N*-glycosylation signal immediately downstream of the CPY leader sequence to track its topology. OM-Psd1p and ER-Psd1p each yielded mature β and *α* subunits as detected by immunoblot of yeast cell extracts demonstrating that self-processing of Psd1p was not impaired by these modifications to its NH_2_-terminus (Fig 2B). All three integrated constructs were similarly expressed relative to each other and over-expressed compared to endogenous Psd1p. Importantly, both OM-Psd1p and ER-Psd1p rescued the ethanolamine auxotrophy of *psd1*Δ*psd2*Δ yeast indicating that these constructs are fully functional and capable of generating levels of PE necessary for cellular growth (Fig 2C).

**Fig 2.**
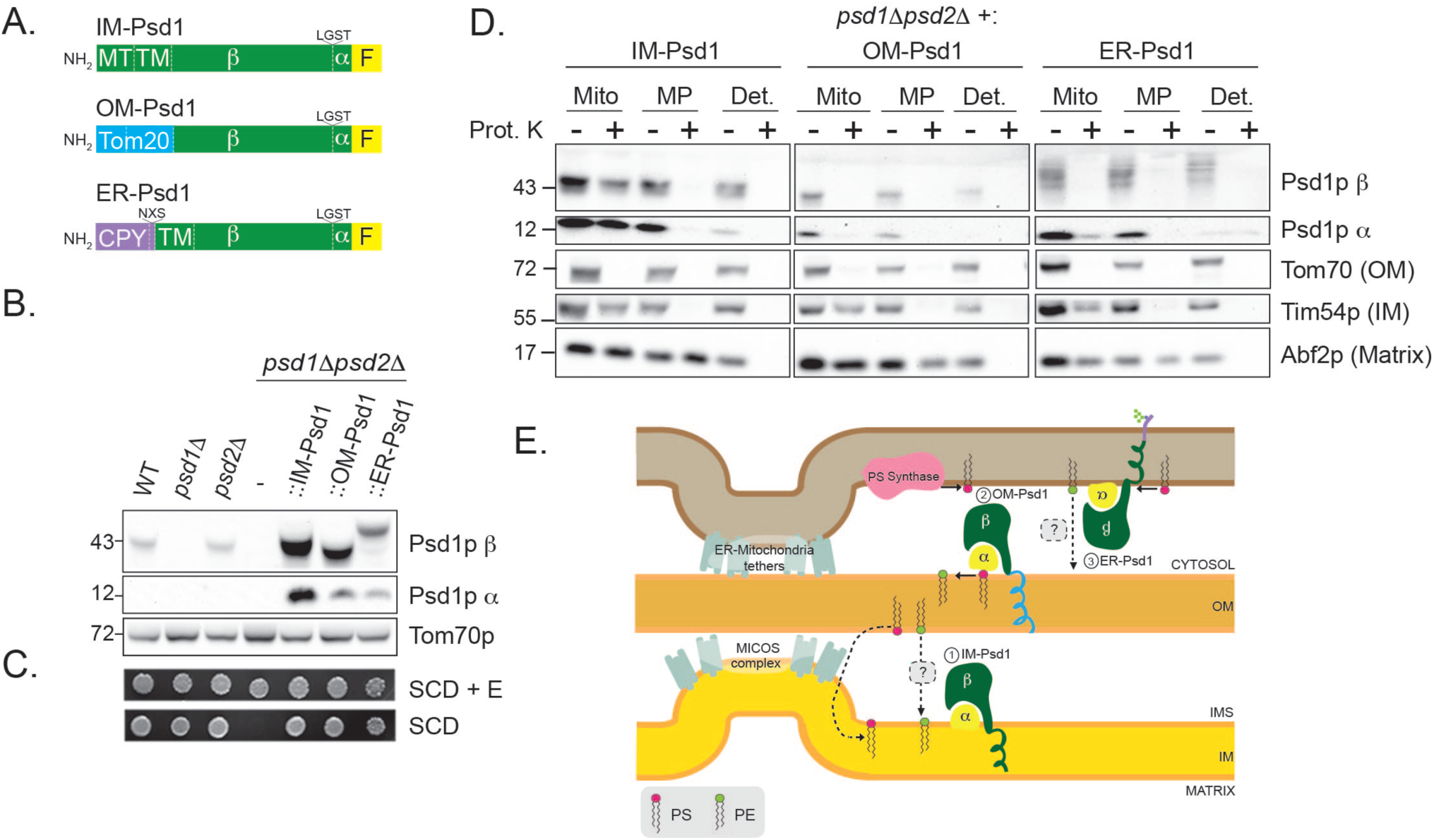
OM-Psd1p and ER-Psd1p constructs are functional and localize to the OM and ER, respectfully. (A) Schematic of IM-Psd1p, OM-Psd1p, and ER-Psd1p. All three constructs contain a 3XFLAG tag at the C-terminus (yellow). The Tom20 residues (1-100) that replace the mitochondrial targeting sequence (MT) and transmembrane (TM) domain of IM-Psd1p (green) are shown for OM-Psd1p (blue), and the carboxypeptidase Y signal sequence (residues 1-37) as well as an NXS motif are indicated for ER-Psd1p (purple). (B) The β and *α* subunits of Psd1p were detected in yeast whole cell extracts of the indicated strains by immunoblot. Tom70p served as a loading control. (C) The indicated strains were spotted onto synthetic complete dextrose (SCD) medium +/- 2mM ethanolamine (+E) and incubated at 30°C for 4 days. (D) Protease protection assay in intact mitochondria (Mito), osmotically ruptured mitochondria (MP), or deoxycholate-solubilized mitochondria (Det.). Following incubation -/+ 100μg proteinase K (Prot. K) for 30 minutes, samples were collected, resolved by SDS-PAGE and immunoblotted for Psd1p (β and α subunits), and the mitochondrial compartment-specific markers Tom70p (OM), Tim54p (IM), and Abf2p (matrix). The exposures of the indicated antibodies were linearly adjusted relative to the set of samples for each strain,and are boxed individually to reflect this. Three biological replicates were performed using two separate batches of isolated mitochondria from each strain. (E) Illustration indicating the topology of (1) IM-Psd1p, (2) OM-Psd1p, and (3) ER-Psd1p.

Previously, localization of ER-Psd1p was established by virtue of its enrichment in the 40,000 × *g* pellet (P40) after subcellular fractionation by gravity centrifugation and its sensitivity to endoglycosidase H, which revealed a mobility shift following SDS-PAGE post-treatment ^29^. To confirm the OM localization of OM-Psd1p, its protease accessibility was determined in intact mitochondria, OM-ruptured mitoplasts, and detergent-solubilized mitochondria and compared to IM-Psd1p. Protease treatment of intact mitochondria expressing IM-Psd1p showed that IM-Psd1p, like the IM control Tim54p, was protected against degradation (Fig 2D). In contrast, similar to the OM control Tom70p, OM-Psd1p was completely degraded in intact mitochondria verifying that it was successfully re-localized to the OM with the bulk of the enzyme facing the cytosol. As expected given the presence of inter-organelle contact sites, a proportion of ER-Psd1p co-fractionated with crude mitochondria (Supplementary Fig 1) and demonstrated protease-sensitivity in intact mitochondria (Fig 2D), a topology that is consistent with its N-glycosylation status (Fig 2E). Thus, a portion of ER-Psd1p is retained in the ER and/or resides in an endosomal compartment that is co-purified with mitochondria.

### OM-Psd1p and ER-Psd1p generate levels of PE in mitochondrial membranes that exceed WT

Next, the lipid content of cellular and mitochondrial membranes was assessed in WT, *psd1*Δ, *psd2*Δ, *psd1*Δ*psd2*Δ, IM-Psd1, OM-Psd1 and ER-Psd1 yeast (Fig 3 and Supplementary Fig 2). The absence of Psd1p resulted in reduced levels of cellular and mitochondrial PE; the combined absence of Psd1p and Psd2p resulted in an additive effect on the steady state abundance of PE (Fig 3A and 3G). In *psd1*Δ and *psd1*Δ*psd2*Δ membranes, the levels of phosphatidylinositol (PI) was increased (Figure 3B and 3H) and CL decreased (Fig 3C and 3I), consistent with previous reports ^9, 28, 33^. A compensatory increase in PS was notably absent in these yeast strains (Fig 3D and 3J). While mitochondrial PE levels were modestly decreased in the absence of Psd2p (Fig 3G), this decrease failed to result in a respiratory growth defect (Fig 1). Combined, these results indicate that Psd2p contributes to the pool of PE associated with mitochondria which is nonetheless unable to functionally replace PE made by Psd1p.

**Fig 3.**
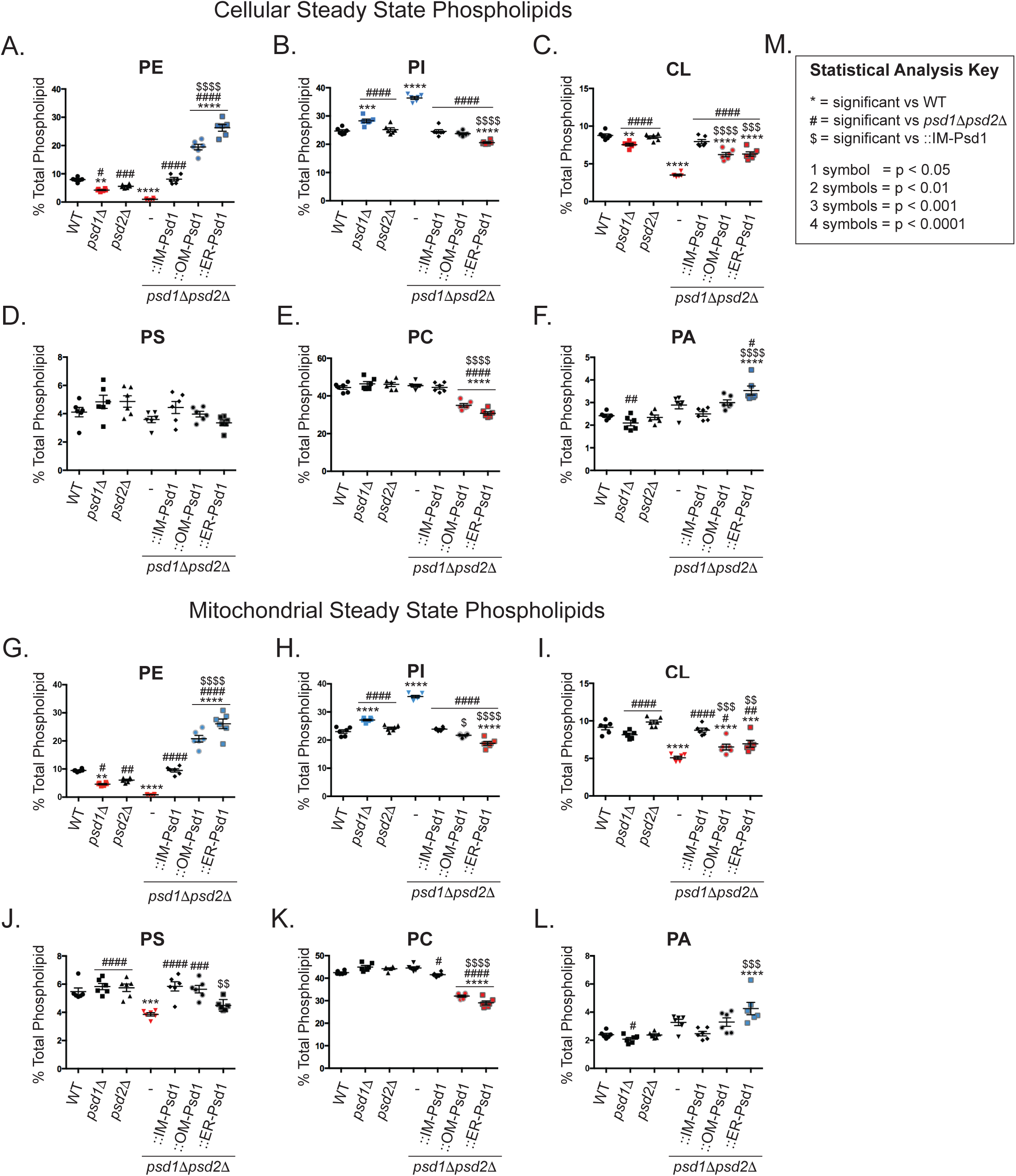
OM-Psd1 and ER-Psd1 contain robust levels of PE in both cellular and mitochondrial membranes. (A-F) Cellular and (G-L) mitochondrial phospholipids from the indicated strains were labeled overnight with ^32^P _i_ and separated by TLC. All graphs show the mean ± S.E.M. for n=6 biological replicates. Significant differences compared to WT were calculated by one-way ANOVA. (M) Key for symbols used for statistical analysis interpretation by one-way ANOVA when comparing samples *versus* WT (*), *psd1*Δ*psd2*Δ (#), or IM-Psd1 ($).

Interestingly, OM-Psd1 and ER-Psd1 yeast contained significantly higher relative amounts of PE than IM-Psd1 in both cellular and mitochondrial membranes (Fig 3A and 3G). As IM-Psd1p, OM-Psd1p, and ER-Psd1p are similarly over-expressed, this suggests that OM-Psd1p and ER-Psd1p, which are either present in compartments where PS is made (ER-Psd1p) or through which PS must traffic to reach IM-Psd1p (OM-Psd1p), have short-circuited normal mitochondrial PE metabolism. In support of this interpretation, the steady state levels of Cho1p (Supplementary Fig 3) and mitochondrial PS (Fig 3J) were increased by OM-Psd1p but not ER-Psd1p, consistent with a mitochondrial trafficking requirement for the former, but not the latter, chimera. Of note, OM-Psd1p and ER-Psd1p both normalized the absolute amount of Cho1p and its phosphorylated pool which were significantly increased in *psd1*Δ*psd2*Δ yeast grown in respiratory conditions (Supplementary Fig 3A, and 3D-F). As Cho1p phosphorylation has been shown to both inhibit enzyme activity and stabilize the polypeptide ^34^, these changes suggest that feedback mechanisms are activated by the severe PE depletion present in *psd1*Δ*psd2*Δ yeast. The relative abundance of phosphatidylcholine (PC) was reduced in OM-Psd1 and ER-Psd1 (Fig 3E and 3K), and although CL levels were significantly increased compared to *psd1*Δ*psd2*Δ mitochondria, they were still lower than in WT (Fig 3C and 3I). The reduced levels of PC in OM-Psd1 and ER-Psd1 is notable as it might have been predicted that an increased production of PE would have resulted in augmented PC synthesis by Pem1p and Pem2p, methyltransferases which reside in the ER and convert PE to PC ^35^. Importantly, the steady state level of Kar2p, the yeast equivalent of the Hsp70 chaperone BiP ^36^, was not increased in OM-Psd1 or ER-Psd1 (Supplementary Fig 3G), demonstrating that their altered membrane compositions did not induce ER stress. In contrast, Kar2p was significantly elevated in both *psd1*Δ*psd2*Δ and *cho1*Δ strains.

Overall, the increased levels of PE detected in OM-Psd1 and ER-Psd1 suggests that both chimeras have increased access to substrate compared to IM-Psd1. This conclusion is bolstered both by the fact that overexpressed IM-Psd1p only restored PE to WT levels (Fig 3A and 3G), as well as results from a prior study that used a plasmid-based overexpression system ^1^. Thus, access to substrate is a major regulatory component that determines Psd1p activity.

A limitation of the radiolabeling-based phospholipid analysis is that it utilized crude mitochondria isolated after physical disruption of intact yeast with glass beads. As such, phospholipid analyses were additionally performed using sucrose gradient purified mitochondria derived from non-radiolabeled yeast cultures. Sucrose purification resulted in mitochondria that were enriched in mitochondrial proteins compared to crude mitochondria (Supplementary Fig 4A). Subsequent phospholipid analyses of the sucrose purified mitochondria provided a profile that was consistent with results derived using crude mitochondria (Fig 3), with one notable exception: the PE levels were roughly equal in OM-Psd1 and ER-Psd1 sucrose purified mitochondria (Supplementary Fig 4E). The reduced amounts of mitochondrial PE in ER-Psd1 following sucrose purification likely reflects their increased purity and provides evidence that PE made in the ER can access mitochondrial membranes. Importantly, there were no dramatic differences in the mitochondrial architectures of strains with elevated or decreased PE levels (Supplementary Fig 4), consistent with previous observations in *psd1*Δ yeast ^28^. Overall, re-routing Psd1p to either the OM or ER results in a robust increase in cellular and mitochondrial PE levels which may or may not reach the mitochondrial IM.

### OM-Psd1 and ER-Psd1 phenocopy the respiratory defect of psd1Δ yeast

We initially evaluated oxidative phosphorylation in these strains by determining their growth on synthetic mediums containing dextrose +/- 2mM ethanolamine, lactate, or ethanol-glycerol +/- 2mM ethanolamine (Fig 4A). Compared to IM-Psd1p, OM-Psd1p and ER-Psd1p only partially improved growth of *psd1*Δ*psd2*Δ yeast on respiratory carbon sources (lactate and ethanol-glycerol). In fact, they supported respiratory growth that was similar to (ER-Psd1p) or slightly better than (OM-Psd1p) *psd1*Δ yeast, but still significantly reduced compared to the WT strain. Interestingly, respiratory growth was slightly better for *psd1*Δ*psd2*Δ yeast expressing OM-Psd1p *versus* ER-Psd1p. The respiratory phenotype of OM-Psd1 and ER-Psd1 yeast suggests that PE made in either the OM or ER cannot functionally replace PE produced in the IM. Indeed, ethanolamine supplementation did not further improve the respiratory growth of either OM-Psd1 or ER-Psd1 yeast (Fig 4A).

**Fig 4.**
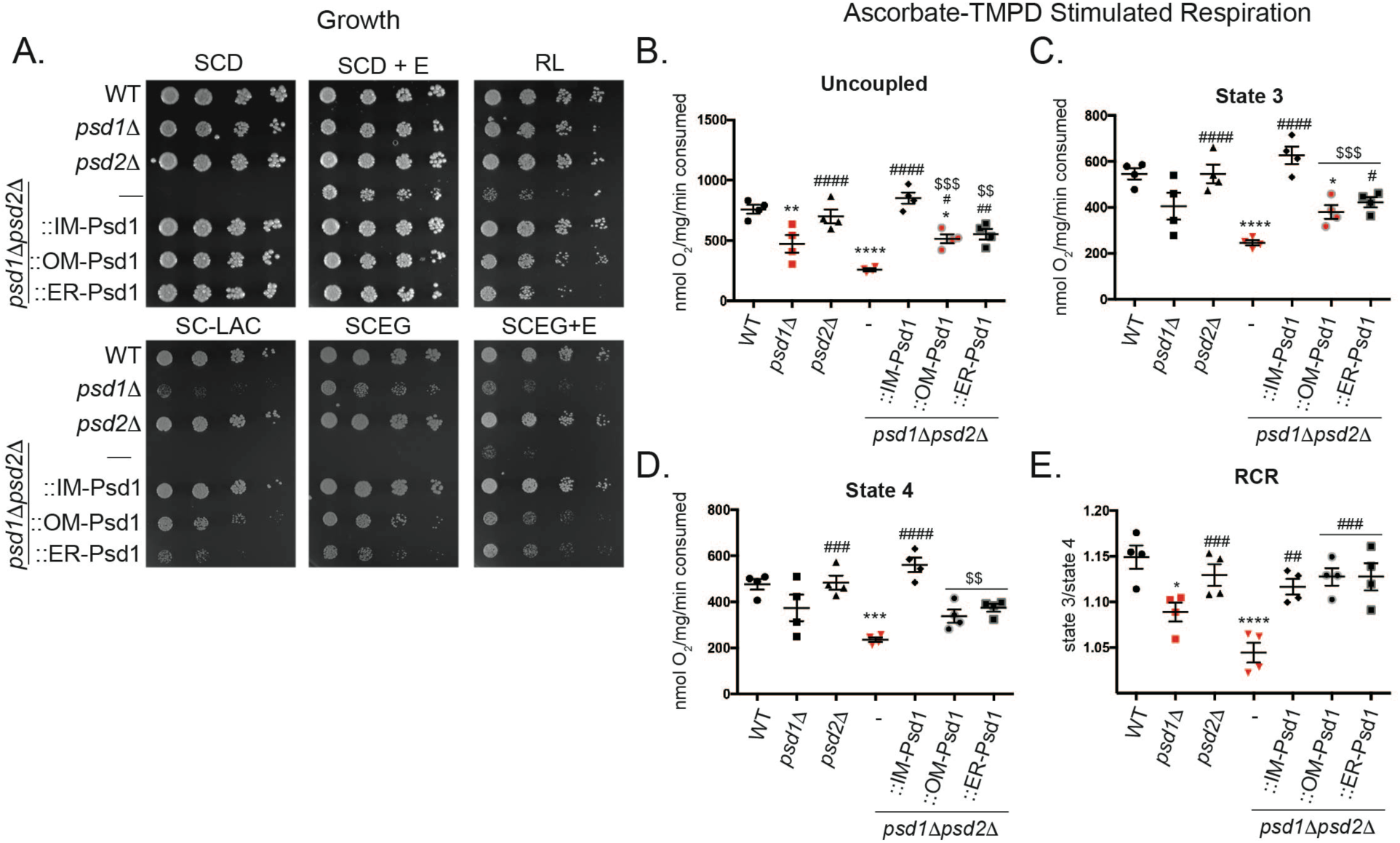
OM-Psd1 and ER-Psd1 OXPHOS function phenocopies *psd1*Δ. (A) The indicated strains were spotted and incubated at 30°C for 2 days on SCD +/- 2mM ethanolamine (+E) and for 5 days on rich lactate (RL), SC lactate (SC-LAC), and SCEG +/- 2mM ethanolamine (+E). (B-E) O_2_ consumption measurements from mitochondria isolated from yeast grown in rich lactate using ascorbate-TMPD as a substrate. (B) The maximal respiratory rate was recorded after the addition of CCCP, (C) state 3 respiration was assessed after addition of ADP, and (D) state 4 respiration was recorded following ADP depletion. (E) The respiratory control ratio (RCR) is calculated by dividing state 3 by state 4 respiratory rates. Analysis *versus* WT (*), *psd1*Δ*psd2*Δ (#), or IM-Psd1 ($) was performed by one-way ANOVA ± S.E.M. for n=4.

To directly assess OXPHOS capacity in these strains, oxygen consumption was monitored in isolated mitochondria using an O_2_ electrode after the addition of ADP and ascorbate tetramethyl-*p*-phenyldiamine (TMPD) which promotes proton pumping by complex IV (Fig 4B-4E). To determine maximal respiratory capacity, carbonyl cyanide *m*-chlorophenyl hydrazine (CCCP) was added after ADP was depleted and mitochondrial respiration had returned from the ADP-stimulated (State 3) to the resting (State 4) respiratory rate. Mitochondria lacking Psd1p, but not Psd2p, had reduced maximal uncoupled respiratory rate compared to WT mitochondria. Even though *psd2*Δ mitochondria consumed O_2_ like WT, the combined absence of Psd1p and Psd2p caused a more severe respiratory defect than was observed when only Psd1p was missing. This indicates that in the absence of Psd1p, PE made by Psd2p has some capacity to compensate by supporting limited complex IV activity. Interestingly, the uncoupled respiratory rate for OM-Psd1 and ER-Psd1 was significantly improved compared to *psd1*Δ*psd2*Δ mitochondria but still impaired relative to IM-Psd1 (Fig 4B). The respiratory control ratio (RCR) is an indication of how well respiration is coupled to ATP synthesis and is calculated by dividing the ADP-stimulated respiration rate by the resting respiration rate (State 3/State 4). RCR ratios were significantly decreased in *psd1*Δ and *psd1*Δ*psd2*Δ mitochondria but perplexingly, OM-Psd1 and ER-Psd1 displayed a ratio similar to WT and IM-Psd1 levels (Fig 4C). For *psd1*Δ*psd2*Δ yeast expressing OM-Psd1p and ER-Psd1p, the normal RCR stems from the fact that they improved State 3 O_2_ consumption more than State 4. Similar to Psd2p in the context of *psd1*Δ yeast, OM-Psd1p and ER-Psd1p significantly improved *psd1*Δ*psd2*Δ respiratory rates to roughly *psd1*Δ levels, indicating that extra-mitochondrial PE can indeed play an important role in enhancing respiration. Combined, these results suggest that defective complex IV function may contribute to the reduced respiratory growth observed when Psd1p is absent from the IM.

### Complex IV function in psd1Δpsd2Δ mitochondria is rescued by ER-Psd1p

When grown in dextrose, *psd1*Δ and *psd1*Δ*psd2*Δ yeast lose their mitochondrial genome, visualized by formation of petite colonies, at a high frequency ^1, 28, 37^. Due to this, we harvested mitochondria from cultures grown in rich lactate to select for growth of respiratory competent cells. As expected, mitochondrial DNA (mtDNA) levels were equivalent between strains in these growth conditions (Fig 5A; ρ^0^ yeast, which are devoid of mtDNA, served as a negative control). Next, isolated complex IV activity measurements were recorded by monitoring the rate of oxidation of reduced cytochrome *c* in mitochondria solubilized in 0.5% *n*-Dodecyl-β-D-maltoside (DDM). Complex IV activity was significantly decreased in *psd1*Δ, *psd1*Δ*psd2*Δ and OM-Psd1 mitochondria but surprisingly, ER-Psd1 retained WT function (Fig 5B). To determine if the different complex IV activities associated with OM-Psd1 and ER-Psd1 mitochondria reflected changes in its expression, we analyzed the steady state amounts of both nuclear and mtDNA encoded subunits of complex IV (Fig 5C). While the levels of the mtDNA-encoded subunit, Cox2p, was significantly decreased in *psd1*Δ*psd2*Δand OM-Psd1 strains, the steady state abundance of the two additional mtDNA-encoded subunits, Cox1p and Cox3p, were not significantly changed and the amount of a constituent encoded in the nucleus, Cox4p, was decreased in OM-Psd1 but not *psd1*Δ*psd2*Δ (Supplementary Fig 5). Further, blue native-PAGE analyses indicated that complex IV assembly into respiratory supercomplexes that consist of a complex III dimer affiliated with either one or two complex IV monomers ^38^, was normal regardless of the absence of *PSD1* or *PSD2*, singly or in combination, as reported by others ^28^, or whether Psd1p was expressed in the IM, OM, or ER (Fig 5D).

**Fig 5.**
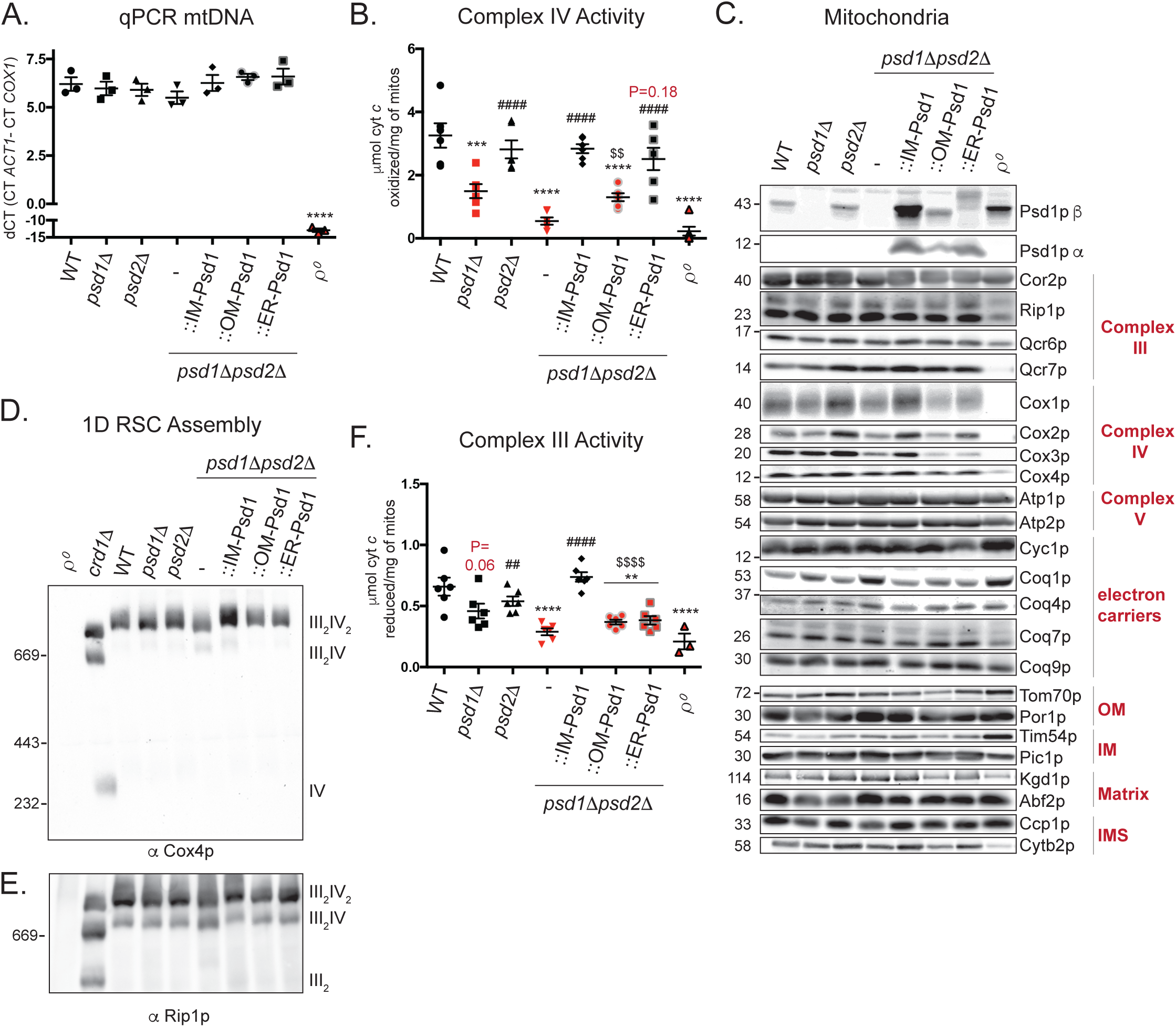
Complex III and IV activities are impaired when Psd1p is absent in the IM. (A) mtDNA was isolated from the indicated strains, normalized, and quantified by qPCR. Analysis was performed by one-way ANOVA ± S.E.M. for n=3. (B) Spectrophotometric assay following the oxidation of cytochrome *c* at 550nM in isolated mitochondria solubilized in 0.5% (w/v) DDM. Analysis *versus* WT (*), *psd1*Δ*psd2*Δ (#), or IM-Psd1 ($) was performed by one-way ANOVA ± S.E.M. for n=6 or n=3 for ρ^0^. P values for decreases that didn’t achieve significance are reported in red and were analyzed by student *t*-test *versus* WT. (C) Mitochondria from the indicated strains were immunoblotted for subunits of complex III (CIII), complex IV (CIV), complex V (CV), and the Coq synthome, cytochrome *c*, and markers of each mitochondrial compartment. (D, E) Blue native-PAGE analysis of respiratory supercomplexes (RSCs) using mitochondrial extracts solubilized in 1.5% (w/v) digitonin. (D) Complex IV assembly was monitored by immunoblot against the nuclear-encoded subunit Cox4p and (E) Complex III assembly was monitored by immunoblot against the nuclear-encoded subunit Rip1p. Mitochondria lacking CL (*crd1*Δ) were used as a positive control for RSC destabilization^53^. (F) Spectrophotometric assay following thereduction of cytochrome *c* at 550nM in isolated mitochondria solubilized in 0.5% (w/v) DDM. Analysis *versus* WT (*), *psd1*Δ*psd2*Δ (#), or IM-Psd1 ($) was performed by one-way ANOVA ± S.E.M for n=6 or n=3 for ρ^0^. P values for decreases that didn’t achieve significance are reported in red and were analyzed by student *t*-test versus WT.

1D RSC Assembly

### Cytochrome bc_1_ complex function is impaired when Psd1p is absent from the IM

The ability of ER-Psd1p, but not OM-Psd1p, to rescue complex IV activity to WT levels was surprising given that neither chimeric construct restored respiratory growth of the *psd1*Δ*psd2*Δ strain to this degree (Fig 4A). Therefore, we postulated that the incomplete respiratory growth rescue of OM-Psd1 and ER-Psd1 could reflect a defect at the level of complex III. Indeed, complex III activity was reduced in *psd1*Δ and significantly decreased in *psd1*Δ*psd2*Δ mitochondrial extracts (Fig 5F). While ER-Psd1p significantly improved complex IV function over that detected in *psd1*Δ*psd2*Δ mitochondria, neither ER-Psd1p or OM-Psd1p supported complex III activity that resembled WT. The reduced complex III activity in *psd1*Δ, *psd1*Δ*psd2*Δ, and *psd1*Δ*psd2*Δ expressing OM-Psd1p or ER-Psd1p did not reflect alterations in the steady state abundance of its subunits (Fig 5C, Cor2p, Rip1p, Qcr6p, and Qcr7p, quantified in Supplementary Fig 5) or its assembly into supercomplexes (Fig 5E) although there was proportionately more of the smaller supercomplex (III_2_IV > III_2_IV_2_) detected in *psd1*Δ*psd2*Δ mitochondrial extracts. Furthermore, the steady state levels of the complex III electron acceptor cytochrome *c* were normal (Fig 5C and Supplementary Fig 6M). Similarly, subunits of the coenzyme Q (CoQ) synthome, a macromolecular complex that catalyzes the synthesis of the complex III electron donor CoQ ^39^, were equal with one exception (Fig 5C and Supplementary Fig 7A). In *psd1*Δ*psd2*Δ mitochondria, Coq1p was increased which could represent an attempt to diminish membrane stress as observed in bacteria ^40^. Moreover, CoQ_6_ supplementation, which is capable of rescuing strains with reduced CoQ biosynthesis ^39^, failed to improve respiratory growth of *psd1*Δ or *psd1*Δ*psd2*Δ yeast (Supplementary Fig 7). Lastly, *psd1*Δ, *psd1*Δ*psd2*Δ, OM-Psd1, and ER-Psd1 respiratory growth was not further impaired in medium lacking *para*-amino benzoic acid (Supplementary Fig 9), a molecule that can be used to produce CoQ by a secondary pathway ^41^. In total, our results demonstrate that CoQ is not limiting for respiratory function in any of these strains. As such, they favor the hypothesis that the impaired complex III activity of *psd1*Δ, *psd1*Δ*psd2*Δ, OM-Psd1, and ER-Psd1 is intrinsic to the multi-subunit holoenzyme itself.

**Fig 6.**
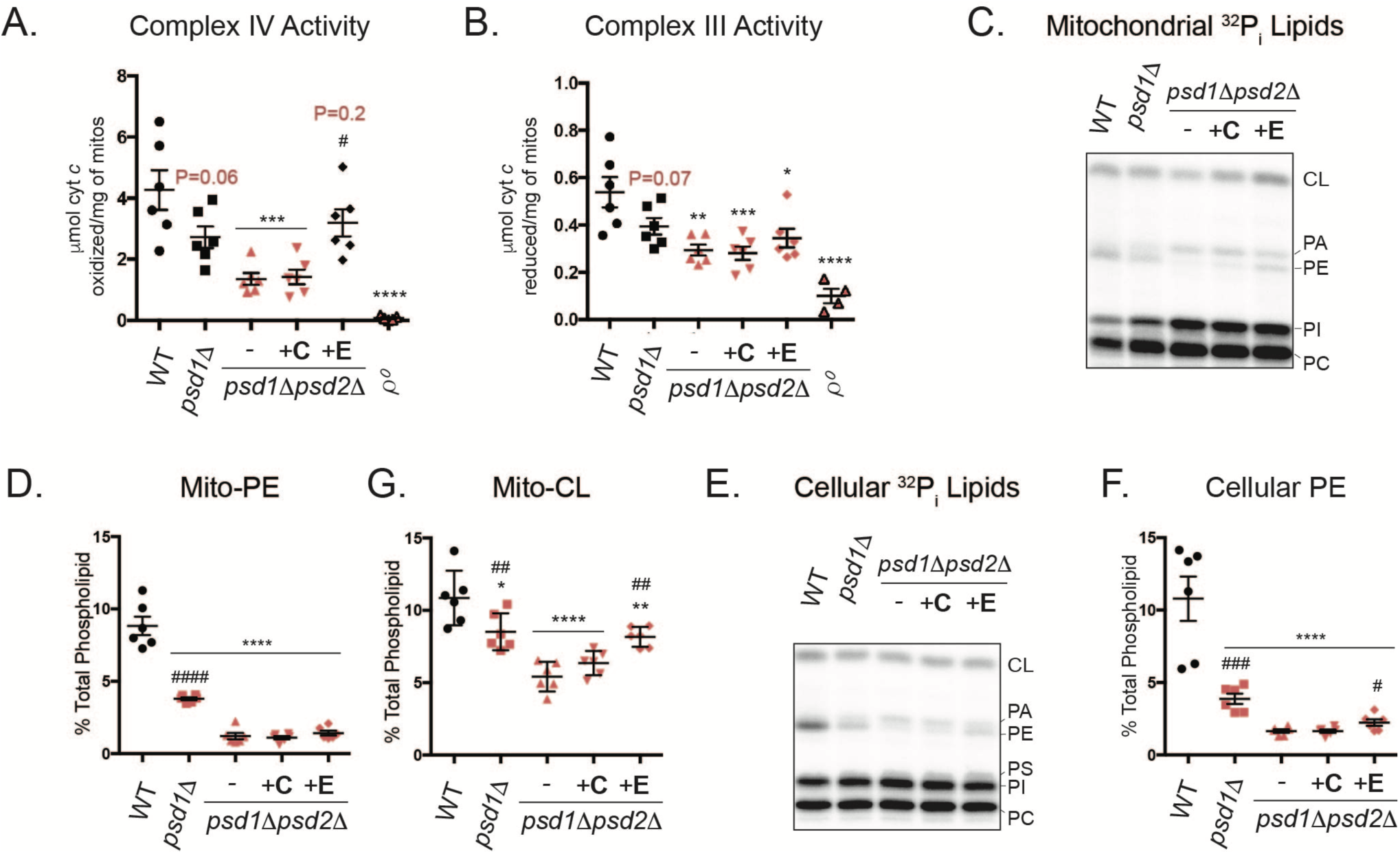
The Kennedy Pathway for PE synthesis can rescue the activity of complex IV but not complex III. Spectrophotometric assay following the (A) oxidation and (B) reduction of cytochrome *c* at 550nM in isolated mitochondria solubilized in 0.5% (w/v) DDM. Analysis *versus* WT (*) or *psd1*Δ*psd2*Δ (#) was performed by one-way ANOVA ± S.E.M. for n=6 or n=4 for ρ^0^. P values for decreases that didn’t achieve significance are reported in red and were analyzed by student *t*-test. (C-G) Mitochondrial phospholipids from the indicated strains were labeled overnight with ^32^P_i_ and separated by TLC. (C) Representative TLC plate for mitochondrial ^32^P_i_ lipids. (D-G) Graphs representing quantitation of (D) mitochondrial PE and (G) CL levels show the mean ± S.E.M. for n=6 biological replicates. Significant differences compared to (*) WT or (#) *psd1*Δ*psd2*Δ were calculated by one-way ANOVA. (E) Representative TLC plate for cellular ^32^P_i_ lipids. (F) Quantitation of cellular PE levels representing the mean ± S.E.M. for n=6 biological replicates. Significant differences compared to (*) WT or (#) *psd1*Δ*psd2*Δ were calculated by student *t*-test.

### PE made by the Kennedy pathway rescues complex IV but not complex III function

As ER-Psd1p restores complex IV, but not complex III, function, we isolated *psd1*Δ*psd2*Δ mitochondria from cultures grown in rich lactate, rich lactate + 2mM choline, and rich lactate +2mM ethanolamine, to evaluate the potential impact of PC and PE generation by the Kennedy pathways on complex IV activity (Fig 6A-B). Similar to ER-Psd1, supplementation of *psd1*Δ*psd2*Δ with ethanolamine, but not choline, restored complex IV (Fig 6A), but not complex III (Fig 6B), function to WT levels. Combined, our results indicate that PE, but not PC, made in the ER by the Kennedy pathway (Fig 6A) or in the endosomal system by either Psd2p or ER-Psd1p (Fig 5B), can significantly rescue the severe complex IV dysfunction that occurs in *psd1*Δ*psd2*Δ yeast. Surprisingly, when we evaluated the phospholipid composition of *psd1*Δ*psd2*Δ yeast supplemented with choline or ethanolamine, we found that the inclusion of ethanolamine did not alter mitochondrial PE levels (Fig 6C-D) and only modestly and yet significantly increased cellular PE abundance (Fig 6E-F), consistent with the slight increases observed in ^1, 33^ but not ^28^. Intriguingly, ethanolamine, but not choline, supplementation significantly increased CL in *psd1*Δ*psd2*Δ yeast to *psd1*Δ levels (Fig 6C and 6G). Significant changes in the abundance of other phospholipid species in *psd1*Δ*psd2*Δ yeast supplemented with either choline or ethanolamine were not observed (Supplementary Fig 10). As such, these results indicate that the Kennedy Pathway for PE but not PC production, is metabolically-linked to CL biosynthesis and/or stability. Moreover, these results suggest that the ability of ethanolamine to improve complex IV activity in *psd1*Δ*psd2*Δ yeast is due to its unanticipated capacity to increase CL levels, a phospholipid known to be important for complex IV function ^42^. These findings further underscore that PE made within the IM is necessary for the full activity of complex III which is otherwise normally expressed, fully assembled, and not limited by the amount of either of its mobile electron carriers.

### Non-enzymatic functions of Psd1p independent of PE biosynthesis are not required for complex III activity

Cho1p mediates the production of PS in the MAM of the ER through conjugation of free serine with CDP-DAG ^18^. In yeast, this feeds into both the Psd1p and Psd2p PS decarboxylation pathways (Fig 7A). We generated deletion strains of Cho1p in the WT and *psd1*Δ*psd2*Δ (labeled *cho1*Δ*1*Δ*2*Δ in Fig 7) genetic backgrounds as a way to deplete mitochondrial PS/PE levels while preserving Psd1p expression. Consistent with previous data, deletion of Cho1p did not affect Psd1p accumulation or maturation ^29^ (Fig 7B) and resulted in an ethanolamine auxotrophy ^43^ (Fig 7C). As anticipated, PS and PE levels were drastically reduced in strains lacking Cho1p (Fig 7D-F). In comparison to *psd1*Δ*psd2*Δ, *cho1*Δ*psd1*Δ*psd2*Δ yeast had a significant increase in CL and PI levels (Fig 7G and 7H), the former of which may be associated with its enhanced respiratory growth (Fig 7K). Deletion of *cho1*Δ in the *psd1*Δ*psd2*Δ background restored PC to WT levels (Fig 7J) but failed to increase the levels of the CL precursor PA (Fig 7I). Growth of *cho1*Δ was decreased compared to WT in YPD and rich lactate medium and similar to *psd1*Δ*psd2*Δ in SCEG supplemented with ethanolamine (Fig 7K). Notably, the activities of complex III (Fig 7L) and IV (Fig 7M) were reduced in the absence of Cho1p. Since Psd1p and essential subunits of complexes III and IV were expressed normally in the absence of Cho1p, singly or in combination with Psd1p and Psd2p (Fig 7N), our combined results directly implicate mitochondrial PE depletion as the cause for the reduced respiratory function of *psd1*Δ, *psd1*Δ*psd2*Δ, and *cho1*Δ yeast.

**Fig 7.**
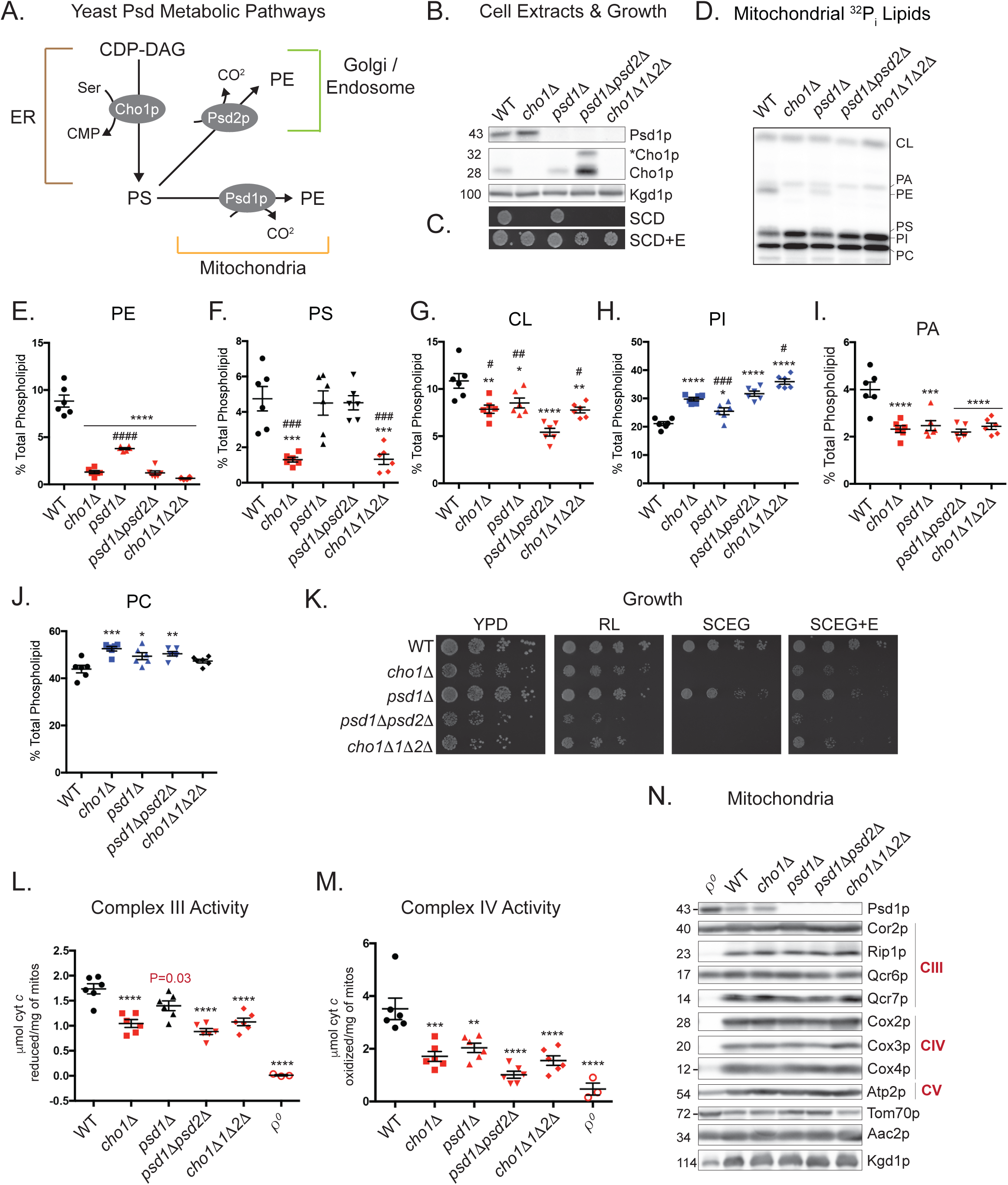
Depletion of PE through deletion of Cho1p impairs complex III and complex IV activities. (A) Metabolic pathways tied to PS synthesis by Cho1p in yeast. (B) Detection of the β subunit of Psd1p and Cho1p expression were verified in yeast whole cell extracts of the indicated strains by immunoblot. Kgd1p served as a loading control. *Cho1p, phosphorylated Cho1p. (C) The indicated strains were spotted onto synthetic complete dextrose (SCD) medium +/- 2mM ethanolamine (+E) and incubated at 30°C for 2 days. (D-J) Mitochondrial phospholipids from the indicated strains were labeled overnight with 32P_i_, separated by TLC, and quantitated by phosphoimaging. All graphs show the mean ± S.E.M. for n=6 biological replicates. Significant differences compared to (*) WT or (#) *psd1*Δ*psd2*Δ were calculated by one-way ANOVA. (K) The indicated strains were spotted and incubated at 30°C for 2 days on YPD and for 3 days on rich lactate (RL), and SCEG +/- 2mM ethanolamine (+E). (L,M) Spectrophotometric assay following the (L) reduction and (M) oxidation of cytochrome *c* at 550nM in isolated mitochondria solubilized in 0.5% (w/v) DDM. Analysis *versus* WT by one-way ANOVA ± S.E.M. for n=6 or n=3 for ρ^0^. (N) Steady state expression of mitochondrial proteins in mitochondria isolated from the indicated strains.

### Glu82 of Qcr7p may coordinate the headgroup of a functionally important PE molecule associated with complex III

PE was identified in crystal structures of the yeast and mammalian cytochrome *bc*_1_ complex in association with the essential mtDNA-encoded catalytic subunit cytochrome *b* (Cob1p) as well as the nuclear-encoded subunit Qcr7p ^30, 31^. Qcr7p is associated with the matrix-facing surface of Cob1p and it is postulated that hydrogen bonding interactions between the headgroup of PE and Glu82 of Qcr7p may help position the complex vertically within the bilayer (Fig 8A). To test the importance of this residue in forming hydrogen bonds with the amine group of PE, we introduced a charge reversal at this position by mutating Glu82 to Arg and also created a strain expressing Asp82 to test the potential effect of shortening the distance of this interaction. Importantly, the amount of Qcr7p^E^82^R^ or Qcr7p^E^82^D^ detected in cell and mitochondrial lysates (Fig 8B and 8C) was similar to WT indicating that neither mutation compromised Qcr7p stability (the Qcr7p^E^82^R^ variant was consistently upshifted compared to WT following SDS-PAGE). Further, Qcr7p^E^82^R^ and Qcr7p^E^82^D^ supported respiratory growth in rich or minimal medium (Fig 8D). Despite being sufficiently functional to promote respiratory growth, complex III activity was significantly decreased for Qcr7p^E^82^R^ and Qcr7p^E^82^D^ to a similar degree as when Psd1p is missing (Fig 8E). Surprisingly, complex IV activity was also decreased in Qcr7p^E^82^R^ but not Qcr7p^E^82^D^ (Fig 8F). The impaired respiratory complex activity for Qcr7p^E^82^R^ and Qcr7p^E^82^D^ was independent of any changes in the steady state amount of their subunits, (Fig 8C). These results provide the first molecular evidence of the functional importance of a conserved PE-binding site identified in the structures of yeast and human complex III. Collectively, these data demonstrate that PE made in the IM by Psd1p is critical to support the intrinsic functionality of complex III and establish one likely mechanism.

**Fig 8.**
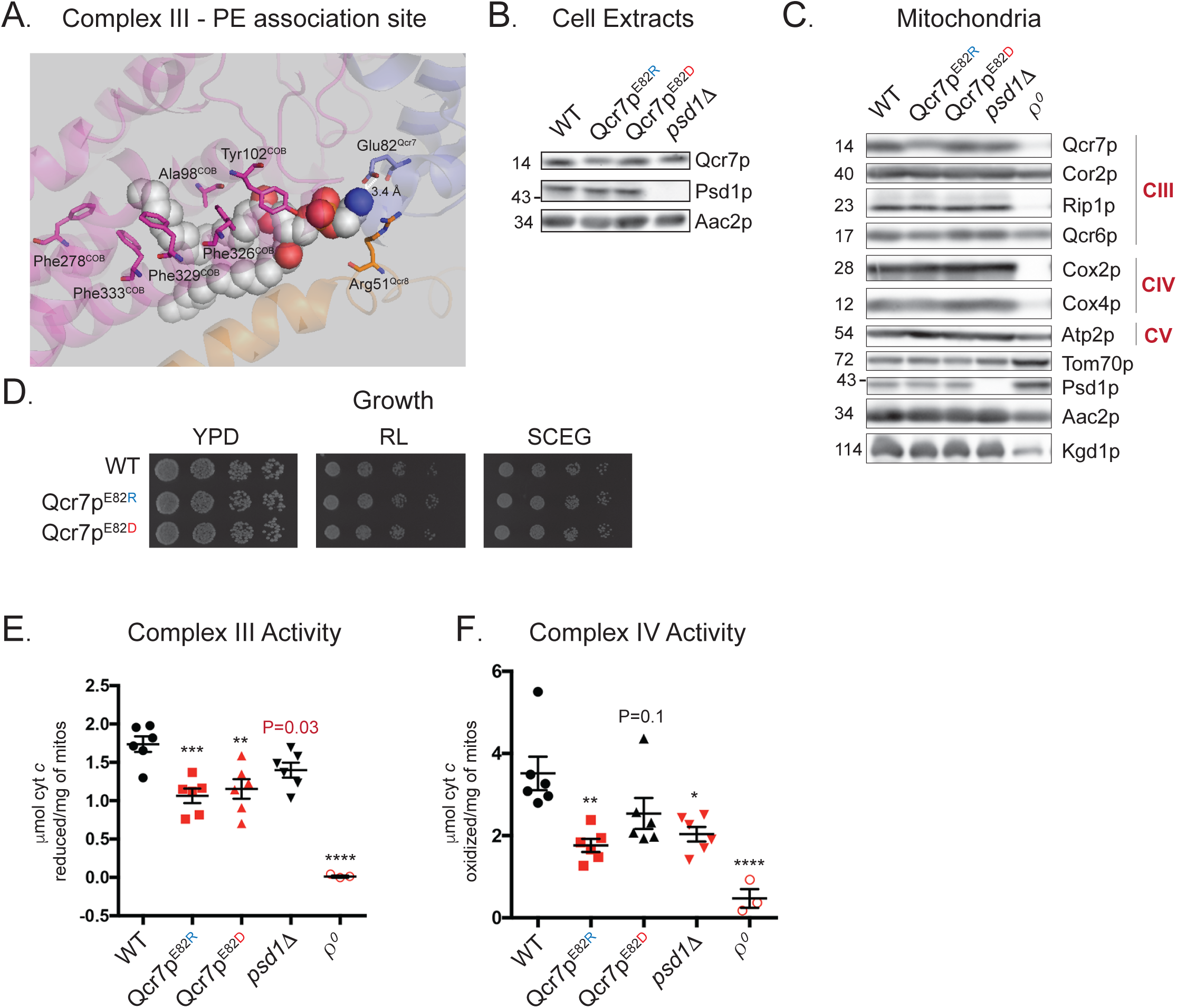
A PE-coordinating residue in Qcr7p is important for complex III activity. (A) The crystal structure of yeast cytochrome *bc*_1_ that modeled associated lipids was downloaded using PDB ID: 1KB9. Using PyMOL, the region containing of the catalytic subunit Cob1p (magenta) near the matrix facing surface was enlarged to demonstrate hydrophobic interactions between this subunit and the acyl chains of PE. Arg51 of Qcr8p (orange) also shows hydrophobic interactions with a carbon atom from the ethanolamine headgroup. Glu82 of Qcr7p (blue) was predicited to form a hydrogen bonding interaction (3.4 Å distance) with the amine group of PE, whose atoms are depicted as spheres *gray*: carbon, *red*: oxygen, and *blue*: nitrogen (hydrogen atoms are not represented). (B) WT and mutant Qcr7p was detected in yeast whole cell extracts on the indicated strains by immunoblot; Aac2p serves as a loading control. (C) Mitochondria from the indicated strains were immunoblotted for subunits of complex III and complex IV as well as markers for the indicated mitochondrial compartments. (D)The indicated strains were spotted and incubated at 30°C for 2 days on YPD and for 3 days on rich lactate (RL) and SCEG. (E, F) Spectrophotometric assay following the (E) reduction or (F) oxidation of cytochrome *c* at 550nm in isolated mitochondria solubilizedin 0.5% (w/v) DDM. Analysis *versus* WT (*) was performed by one-way ANOVA ± S.E.M. for n=6 or n=3 for ρ^0^. P values for decreases that didn’t achieve significance are reported and were evaluated by student t-test *versus* WT.

## DISCUSSION

PE levels have been shown to be altered in models of the neurodegenerative disorders Parkinson’s disease ^44^ and Alzheimer’s disease ^45^, and recent evidence has implicated the turnover of mammalian Pisd as an important regulatory mechanism that diminishes the proliferation of breast cancer cells ^46^. Thus, regulation of the Psd pathway is not only important for mitochondrial function but also organismal health and physiology.

Using an assortment of biochemical assays that measure both the combined and individual activities of the respiratory chain complexes, we sought to clarify the respiratory defect associated with *psd1*Δ yeast and whether or not PE made by the Kennedy pathway could rescue it. Two different groups using the same yeast strain background have reported that supplemental ethanolamine rescues ^28^ or fails to rescue ^11^ respiratory function of *psd1*Δ yeast. Given these conflicting results, we first tested the ability of ethanolamine supplementation to improve the respiratory growth phenotype of yeast lacking Psd1p. While we observed that growth of *psd1*Δ*psd2*Δ could be rescued on respiratory medium supplemented with ethanolamine, growth after 3 days was delayed and impaired similarly to *psd1*Δ yeast. Perplexingly, supplementation of *psd1*Δ*psd2*Δ with ethanolamine to generate PE through the Kennedy pathway did not significantly increase mitochondrial PE levels and instead resulted in a significant, albeit modest, increase in CL (Fig 6C-F), which nonetheless failed to completely restore respiratory growth. Importantly, the inability of ethanolamine to fully restore *psd1*Δ respiratory growth was verified in four distinct yeast strain backgrounds suggesting that genetic differences are not the source for the disparate results reported previously ^1, 11, 27, 28^. This notion is further supported by the fact that two groups that each utilized *psd1*Δ derived from the same parental strain came to opposite conclusions regarding the rescuing capacity of PE made by the Kennedy pathway ^11, 28^.

Interestingly, lyso-PE supplementation (PE containing only one acyl chain) can rescue *psd1*Δ respiratory growth but acts via a separate pathway than the Kennedy pathway ^27^. Further, lyso-PE has also been demonstrated to reverse the mitochondrial defects associated with RNAi silencing of mammalian *Pisd* in cell culture ^10^, and can increase the proliferation rate of the MCF7-RAS breast cancer cell line when Pisd levels are depleted ^46^. Thus, the different ER-resident PE producing pathways appear to differ with respect to their ability to compensate for the absence of the mitochondrial Psd pathway suggesting that pathway-specific mitochondrial import mechanisms for PE may exist.

Our oxidative phosphorylation analyses demonstrated that both OM-Psd1 and ER-Psd1 phenocopy the respiratory defect of *psd1*Δ mitochondria. This implies that the IMS is the greatest barrier preventing PE import into the inner membrane. Moreover, we have identified a key oxidative phosphorylation component that is uniquely dependent on PE made by Psd1p in the IM: respiratory complex III. Specifically, we demonstrated that while PE synthesized in the endosomal system–by ER-Psd1p, Psd2p, or the Kennedy pathway– can rescue the impaired complex IV activity that occurs in the absence of Psd1p in the IM, it fails to do so for complex III (Fig 5F and Fig 6B). Since complex III was impaired in yeast deficient in mitochondrial PS and PE that contained a normal amount of fully processed Psd1p (*cho1*Δ, Fig 7K), our results demonstrate that the complex III functional defect results from an insufficient supply of PE and not from a non-PE related activity of Psd1p.

What is the basis for the capacity of extra-mitochondrially produced PE to rescue complex IV but not complex III function? As both complex III and complex IV are IM residents, one possibility is that these respiratory complexes require different levels of PE within the inner membrane to function appropriately. If true, this would imply that the relative levels of PE in the IM in the strains analyzed are equivalent between WT, *psd2*Δ, and IM-Psd1, similarly depleted in ER-Psd1 and *psd1*Δ, less than *psd1*Δ in OM-Psd1, and almost completely absent in *psd1*Δ*psd2*Δ. Unfortunately, while it is clear that OM-Psd1 and ER-Psd1 mitochondria contain robust levels of PE, we have yet to determine its steady state distribution between the outer and inner membranes. Another possible explanation for the different dependence of complex III and IV function on IM-produced PE could reflect the nature of the PE made in the mitochondrion versus elsewhere. Evidence that polyunsaturated 38:4 and 38:5 PE species are enriched in the mitochondrial inner membrane of HeLa cells has been previously reported ^47^. While both the Kennedy and Psd pathways have the capacity to produce all diacyl-PE species, the Kennedy pathway preferentially synthesizes mono 34:2 or di-unsaturated 36:3 PE and Pisd mainly generates polyunsaturated 38:4 or 38:5 PE ^48^. However, *S. cerevisiae* is unable to synthesize complex unsaturated fatty acids and the PE acyl chain pattern is the same between whole cells and isolated mitochondria ^49, 50^. It is interesting to note that, even though the Kennedy and Psd pathways both share a preference for generating both mono-and di-unsaturated PE in yeast, there is a 10% enrichment of 34:2 PE that contains oleic and palmitoleic acid in crude mitochondria versus ER-derived microsomes ^49^. Another intriguing possibility is that the synthesis of PE by Psd1p in the IM is somehow directly coupled to the incorporation of this lipid into partially or fully assembled complex III. Recently, a subunit of complex IV was found to associate with the MICOS subunit, Mic19, in mammalian cells by EM tomography and immunoprecipitation ^51^. As the MICOS complex was suggested to work in concert with Ups2p to regulate mitochondrial PS/PE metabolism in yeast ^26^, contact sites between the OM and IM could potentially facilitate the transport of ER-derived PE to complex IV more directly than it can to complex III.

Intriguingly, the improved complex IV functionality provided by ethanolamine correlated best with an increased abundance of CL and not PE (Fig 6C-F), a phenomenon that has been observed previously ^1, 28^, suggesting that this rescue is actually CL-dependent. The importance of CL in the assembly and function of the respiratory complexes and supercomplexes is well documented ^42, 52, 53^. Our results strongly indicate that when Psd1p is missing or mis-localized, a minimal threshold of PE is not achieved within the IM that is necessary to accumulate normal levels of CL and promote the full activity of the OXPHOS machinery. Although it is presently unclear how the Kennedy Pathway for PE production promotes CL accumulation, it is known that the Ups1p and Ups2p lipid trafficking proteins have an inverse relationship with respect to CL/PE metabolism suggesting that this may be linked to PS/PA trafficking into the IM ^3, 54, 55^. Depletion of PS in the *psd1*Δ*psd2*Δ background restored CL levels to that of *psd1*Δ and *cho1*Δ which were still comparatively reduced *vs* WT (Fig 7G). This increase in CL also coincided with improved growth for *cho1*Δ*psd1*Δ*psd2*Δ yeast (Fig 7K). It is possible that in the absence of PS, metabolic pathways that either promote PA formation in the ER ^56^ or promote PA import to the IM ^57, 58^ are stimulated. Moving forward, it will be important to distinguish between these non-mutually exclusive models.

Another outstanding question raised by our study is exactly how PE made by Psd1p in the IM promotes complex III function. Since *psd1*Δ*psd2*Δ yeast contain normal amounts of cytochrome *c* and CoQ_6_ is not limiting, the underlying respiratory defect is intrinsic to complex III. We demonstrated that mutations in a residue of Qcr7p predicted to bind PE impaired complex III activity (Fig 8E) establishing a likely mechanistic basis for the requirement of PE for its function. To our knowledge, this is the first molecular evidence demonstrating the functional importance of a specific interaction of PE with complex III, which until now had only been postulated from crystal structures of yeast complex III and human respiratory supercomplexes ^30, 31^. A second PE is found adjacent to the dimer interface of this complex; as such, the acyl chains of PE are thought to interact with both monomers and assist in dimer formation. Additionally, the acyl chains of PE at the dimer interface could potentially promote quinol-quinone exchange at the Q_i_ and Q_0_ sites. More broadly, during quinol-quinone exchange, sidechain movement of Cob1p is thought to be necessary to transfer protons from His202 to ubiquinone, and perhaps depletion of PE hinders this movement and delays proton flux ^59^.

Further, electron-transfer reactions occur within microsecond-to-millisecond time scales and electron-tunneling between complex III monomers is thought to maximize the efficiency of coupled redox reactions between monomers ^60^. The generation of the free radical semiquinone during the Q cycle performed by complex III is proposed to be diminished by a mechanism that delays quinol oxidation in the Q_o_ site when electron transfer is pharmacologically impaired ^61^. If depletion of PE or mutagenesis of residues that interact with this lipid diminish the efficiency of electron transfer between complex III monomers or between complex III and its substrates, this could result in reduced complex III function as a means to prevent superoxide production. These structural observations will guide future efforts to determine the role(s) of PE as it relates to complex III activity.

## MATERIALS AND METHODS

### Yeast strains and growth conditions

All yeast strains used in this study are listed in Table 1 and were derived from GA74-1A unless otherwise noted. Deletion strains were generated by PCR-mediated gene replacement of the entire reading frame as previously described ^52^. Psd1p containing a COOH-terminal 3XFLAG tag subcloned into pRS305 has been described ^17, 29^. To re-direct Psd1p to the mitochondrial OM, the first 100 amino acids of Psd1p, encompassing its mitochondrial targeting sequence and transmembrane domain, were replaced by the equivalent domains (amino acids 1-34) of the single-pass OM resident protein, Tom20p. ER-Psd1p, which is directed to the secretory pathway, was generated by replacing the first 57 amino acids of Psd1p, encompassing its mitochondrial targeting signal (MTS), with the N-terminal signal sequence (amino acids 1-23) of carboxypeptidase Y, as previously described ^29^. Additionally, the ER-Psd1p construct contains an NXS *N*-glycosylation signal immediately downstream of the CPY leader sequence to track its topology. The IM-Psd1, OM-Psd1, and ER-Psd1 constructs, which all contained the C-terminal 3XFLAG tag, were subcloned into the pRS305 plasmid, linearized, and integrated into the *LEU2* locus in the *psd1*Δ*psd2*Δ background. Clones were selected on synthetic dropout medium (0.17% (w/v) yeast nitrogen base, 0.5% (w/v) ammonium sulfate, 0.2% (w/v) dropout mixture synthetic-leu, 2% (w/v) dextrose) and verified by immunoblot.

**TABLE 1:** Strains used in this work

| TABLE 1 - Strains used in this work |  |  |
| --- | --- | --- |
| Strain | Genotype | Source |
| **Parental** |  |  |
| GA74-1A | *MATa, his3-11, 15, leu2, ura3, trp1, ade8 [rho+, mit+]* | Carla Koehler |
| **Deletions** |  |  |
| *psd1Δ* | *MATa, psd1Δ::HIS3MX6, leu2, ura3, trp1, ade8 [rho+, mit+]* | Ref. 29 |
| *psd2Δ* | *MATa, psd2Δ::HIS3MX6, leu2, ura3, trp1, ade8 [rho+, mit+]* | Ref. 29 |
| *psd1Δpsd2Δ* | *MATa, psd2Δ::HIS3MX6, leu2, ura3, psd1Δ::TRP1, ade8 [rho+, mit+]* | Ref. 29 |
| *crd1Δ* | *MATa, his3-11, 15, leu2, ura3, crd1Δ::TRP1, ade8 [rho+, mit+]* | Ref. 67 |
| $\rho^0$ | *MATa, his3-11, 15, leu2, ura3, trp1, ade8 [rho-, mit+]* | This study |
| *cho1Δ* | *MATa, cho1Δ::HIS3MX6, leu2, ura3, trp1, ade8 [rho+, mit+]* | Ref. 29 |
| **Integrations** |  |  |
| *psd1Δpsd2Δ::IM-Psd1* | *MATa, psd2Δ::HIS3MX6, Psd3XFLAG::LEU2, ura3, psd1Δ::TRP1, ade8 [rho+, mit+]* | Ref. 29 |
| *psd1Δpsd2Δ::OM-Psd1* | *MATa, psd2Δ::HIS3MX6, Tom20-Psd3XFLAG::LEU2, ura3, psd1Δ::TRP1, ade8 [rho+, mit+]* | This study |
| *psd1Δpsd2Δ::ER-Psd1* | *MATa, psd2Δ::HIS3MX6, CPY\*mPsd3XFLAG::LEU2, ura3, psd1Δ::TRP1, ade8 [rho+, mit+]* | Ref. 29 |
| **HI-CRISPR Edited** |  |  |
| *cho1Δpsd1Δpsd2Δ* | *MATa, psd2Δ::HIS3MX6, leu2, ura3, psd1Δ::TRP1, \*cho1 HI-CRISPR ade8 [rho+, mit+]* | This study |
| *Qcr7p E82R* | *MATa, his3-11, 15, leu2, ura3, trp1, ade8 [rho+, mit+] \*Qcr7p<sup>E82R</sup> HI-CRISPR* | This study |
| *Qcr7p E82D* | *MATa, his3-11, 15, leu2, ura3, trp1, ade8 [rho+, mit+] \*Qcr7p<sup>E82D</sup> HI-CRISPR* | This study |
| **Parental/Deletion** |  |  |
| W303 | *MATa, his3-11, ura3-1, 15, trp1-1, ade2-1, can1-100 [rho+, mit+]* | Cathy Clarke |
| *psd1Δ* W303 | *MATa, psd1Δ::HIS3MX6, ura3-1, 15, trp1-1, ade2-1, can1-100* | This study |
| D2730-10B | *MATa [rho+, mit+]* | Carla Koehler |
| *psd1Δ* D2730-10B | *MATa, psd1Δ::KANMX6, [rho+, mit+]* | This study |
| BY4741 | *MATa, his3-1, leu2-0, met15-0, ura3-0 [rho+, mit+]* | Euroscarf |
| *psd1Δ* BY4741 | *MATa, psd1Δ::KANMX4, his3-1, leu2-0, met15-0, ura3-0 [rho+, mit+]* | Euroscarf |

For strains that were genetically modified by homology-integrated clustered regulatory interspaced short palindromic repeats (CRISPR)-Cas (HI-CRISPR) ^62^, CRISPR-Cas9 gene blocks were designed to target *CHO1* or *QCR7* and cloned into the pCRCT plasmid ^62^, as previously described ^17^. The spacer sequences (including the CRISPR-Cas9 target (20-bp) and the homology repair template (50-bp) homology arms on both sides (total 100-bp) flanking the Cas9 recognition sequence) were ordered as gBlocks (Integrated DNA Technologies). Specifically, *CHO1* knockout was achieved by incorporating an 8-bp deletion within the homology repair template to induce a frameshift mutation by removal of nucleotides 38-45 downstream of the start ATG sequence of the *CHO1* open reading frame (ORF). The CRISPR construct was designed to target the protospacer adjacent motif (PAM) sequence encoded by nucleotides 40-42 on the reverse strand of the *CHO1* ORF. Single point mutations were introduced into the *QCR7* ORF at positions 244-246 downstream of the ATG start site to mutate Glu82 (GAG) to Asp82 (GAC) or Arg82 (AGA) through the design of homology repair templates encoding these mutations. Specificity of gene block integration into this locus was accomplished by targeting the PAM sequence encoded by nucleotides 209-211 on the reverse strand of the *QCR7* ORF. Both *QCR7* gene blocks encoded for silent alanine mutations at position 208-210 to mutate the PAM sequence and prevent re-cleavage by Cas9 after homology-directed repair, and position 214-216 to prevent hairpin formation of gBlocks. Additionally, the homology arms adjacent to the PAM sequence were extended to 125-bp on each side for these constructs. Integration of gene blocks encoding for these mutations were verified by sequencing of yeast genomic DNA using primers specific for *QCR7*.

Yeast were grown on YPD (1% (w/v) yeast extract, 2% (w/v) tryptone, 2% (w/v) dextose) plates. To assess the function of the assorted re-directed Psd1p constructs, overnight cultures grown in synthetic complete dextrose (SCD; 0.17% (w/v) yeast nitrogen base, 0.5% (w/v) ammonium sulfate, 0.2% (w/v) complete amino acid mixture, 2% (w/v) dextrose) supplemented with 2mM ethanolamine hydrochloride were spotted on SCD plates in the absence or presence of 2mM ethanolamine hydrochloride, or spotted onto rich lactate (1% (w/v) yeast extract, 2% (w/v) tryptone, 0.05% (w/v) dextrose, 2% (v/v) lactic acid, 3.4mM CaCl_2_-2H_2_O, 8.5mM NaCl, 2.95mM MgCl_2_-6H_2_O, 7.35mM KH_2_PO_4_, 18.7mM NH_4_Cl, pH 5.5), or synthetic complete ethanol glycerol (SCEG; 0.17% (w/v) yeast nitrogen base, 0.5% (w/v) ammonium sulfate, 0.2% (w/v) complete amino acid mixture, 1% (v/v) ethanol, 3% (v/v) glycerol) or synthetic complete lactate (SC-LAC; 0.17% (w/v) yeast nitrogen base, 0.5% (w/v) ammonium sulfate, 0.2% (w/v) complete amino acid mixture, 0.05% (w/v) dextrose, 2% (v/v) lactic acid, 3.4mM CaCl_2_-2H_2_O, 8.5mM NaCl, 2.95mM MgCl_2_-6H_2_O, 7.35mM KH_2_PO_4_, 18.7mM NH_4_Cl, pH 5.5) in the absence or presence of 2mM ethanolamine hydrochloride to test respiratory growth.

For CoQ_6_ supplementation experiments, starter cultures were grown in SCD + 2mM ethanolamine overnight and diluted to 0.025 OD_600_ in 500μL of either SCEG, SCEG + 2mM ethanolamine, SCEG + 2μM CoQ_6_ (Avanti Polar Lipids, Inc), SCEG + 2μM CoQ_6_ + 2mM ethanolamine, SCEG + 10μM CoQ_6_, or SCEG + 10μM CoQ_6_ + 2mM ethanolamine in duplicate in a 48 well plate. OD_600_ measurements were then recorded every 30 minutes for a period of 48 hours at 30°C using a Tecan Infinite 200 Pro instrument. For experiments designed to test the importance of the pABA pathway for cell growth, synthetic media lacking pABA was utilized for liquid and solid growth of yeast cells. -pABA media consisted of 790 mg/mL CSM Mixture Complete (Formedium LTD) and 6.9g/L yeast nitrogen base lacking amino acids and *para*-amino benzoic acid (Formedium LTD). For +pABA media, the same mixture was used but included the addition of 100μM *para*-amino benzoic acid (Research Products International, Inc) dissolved in water and sterile filtered using a 0.20μM filter (Corning, Inc). Individual colonies of yeast were used to inoculate starter cultures in -pABA medium containing 2% (w/v) glucose and 2mM ethanolamine. Solid growth of yeast on agar plates was evaluated in -/+pABA medium containing 1% (v/v) ethanol and 3% (v/v) glycerol -/+ 2mM ethanolamine.

### Mitochondrial Isolation

Isolation of crude mitochondria was performed as previously described ^63^. Yeast were selected on rich lactate plates and grown on rich lactate media to prevent the loss of mitochondrial DNA. To improve growth on rich lactate, *psd1*Δ*psd2*Δ yeast were grown in the presence of 2mM choline prior to harvesting mitochondria (except where indicated in Fig 6A,B and in Fig 7). Purification of mitochondria by sucrose step gradient was performed as previously described ^29^.

### Electron microscopy

Cells were grown in rich lactate medium and harvested at mid-log phase by centrifugation. Cells were fixed in 3% glutaraldehyde contained in 0.1 M sodium cacodylate, pH 7.4, 5 mM CaCl_2_, 5 mM MgCl_2_, and 2.5% (w/v) sucrose for 1 hr at room temperature with gentle agitation, spheroplasted, embedded in 2% ultra-low temperature agarose (prepared in water), cooled, and subsequently cut into small pieces (∼ 1 mm^3^) as previously described ^52^. The cells were then post-fixed in 1% OsO_4_, 1% potassium ferrocyanide contained in 0.1 M sodium cacodylate, 5 mM CaCl_2_, pH 7.4, for 30 min at room temperature. The blocks were washed thoroughly four times with double distilled H_2_O, 10 min total, transferred to 1% thiocarbohydrazide at room temperature for 3 min, washed in double distilled H_2_O (four times, 1 min each), and transferred to 1% OsO_4_, 1% potassium ferrocyanide in 0.1 M sodium cacodylate, pH 7.4, for an additional 3 min at room temperature. The cells were washed four times with double distilled H_2_O (15 min total), stained *en bloc* in Kellenberger’s uranyl acetate for 2 hr to overnight, dehydrated through a graded series of ethanol, and subsequently embedded in Spurr resin. Sections were cut on a Reichert Ultracut T ultramicrotome, post-stained with uranyl acetate and lead citrate, and observed on an FEI Tecnai 12 transmission electron microscope at 100 kV. Images were recorded with a Soft Imaging System Megaview III digital camera, and figures were assembled in Adobe Photoshop with only linear adjustments in contrast and brightness.

### mtDNA quantitation

DNA was extracted as described ^64^. In brief, yeast cells were grown for 2 days in rich lactate and the collected cell pellets were vortexed at level 10 for 3 minutes with 200μL breaking buffer (2% (v/v) Triton X-100, 1% (v/v) SDS, 100mM NaCl, 10mM Tris pH8.0, 1mM EDTA pH 8.0), 0.3g glass beads, and 200μL phenol/chloroform/isoamyl alcohol at room temperature. The solution was neutralized with the addition of 200μL of Tris-EDTA (TE) buffer pH 8.0 and phases were separated by centrifugation at 21,000 × *g* for 5 minutes. The aqueous phase was collected and DNA was precipitated by the addition of 100% ethanol and collected in the pellet after centrifugation at 21,000 × *g* for 3 minutes. The pellets were resuspended in 400μL TE buffer, pH 8.0 and RNA was digested with the addition of 3 μL of 10mg/mL RNAse A and incubation at 37°C for 5 minutes before addition of 10 µL of 4M Ammonium acetate and 1 mL 100% ethanol. DNA pellets were recovered by centrifugation at 21,000 × *g* for 3 minutes, dried, and resuspended in 30 µL TE buffer pH 8.0. DNA was stored at −80°C until ready for use, quantitated, and a portion diluted to 10ng/μL to be used as template in the qPCR reaction. The FastStart Universal SYBR Green Master Rox (Roche) was used for qPCR performed according to the manufacturer’s instructions. 50ng of genomic DNA was used as a template and the following primers were used at 100nM concentration in a 20µL reaction: *COX1* forward (5′- CTACAGATACAGCATTTCCAAGA-3′), *COX1* reverse (5′- GTGCCTGAATAGATGATAATGGT- 3′), *ACT1* forward (5′- GTATGTGTAAAGCCGGTTTTG-3′), and *ACT1* reverse (5′- CATGATACCTTGGTGTCTTGG -3′). The reactions were performed in technical duplicate with three biological replicates. After completion of thermocycling in a QuantStudio 6 Flex Real-Time PCR System (Thermo Fisher), melting-curve data was collected to verify PCR specificity. The absence of primer dimers and the Ct value difference between the nuclear (*ACT1*) and mitochondrial (*COX1*) target were computed as a measure of the mitochondrial DNA copy number relative to the nuclear genome.

### Mitochondrial respiration measurements

Mitochondrial oxygen consumption was measured using a Clark-type oxygen electrode in a magnetically stirred, thermostatically controlled 1.5 mL chamber at 25°C (Oxytherm; Hansatech) as previously described ^65^ with some variations. 100μg of mitochondria were resuspended in 0.25M sucrose, 0.25mg/mL BSA, 20mM KCl, 20mM Tris-Cl, 0.5mM EDTA, 4mM KH_2_PO_4_, and 3mM MgCl_2_, pH 7.2. After addition of 1mM ascorbate + 0.3mM TMPD, state 2 rate was monitored for approximately 30 sec. State 3 respiration was initiated by addition of 50μM ADP. After state 4 rate was measured, 10μM CCCP was added to induce uncoupled respiration, and the rate was followed for either 2 minutes or until oxygen level reached zero.

### Complex III and IV Activity Measurements

Complex III and IV activities were measured as described ^66, 67^. To measure complex III activity, 25μg of mitochondria solubilized in 0.5% (w/v) *n*-dodecyl-β-D-maltoside were added to reaction buffer (50mM KP_i_, 2mM EDTA, pH 7.4) with 0.008% (w/v) horse heart cytochrome *c* and 1mM KCN. The reaction was started by adding 100μM decylubiquinol, and the reduction of cytochrome *c* followed at 550nM. Complex IV activity was initiated by adding 5μg of solubilized mitochondria to reaction buffer with 0.008% ferrocytochrome *c* and measured by recording cytochrome *c* oxidation at 550nm.

### Antibodies

Most antibodies used in this study were generated by our laboratory or in the laboratories of J. Schatz (University of Basel, Basel, Switzerland) or C. Koehler (UCLA) and have been described previously ^3, 63, 65, 67-69^. Other antibodies used were mouse anti-Sec62p (kind gift of Dr. David Meyers (UCLA)), mouse anti-FLAG (clone M2, Sigma), mouse anti-Dpm1p (113686, Abcam), rabbit anti-Qcr7p ^70^, rabbit antisera reactive to Coq1p ^71^, Coq4p ^72^, Coq7p ^73^, or Coq9p ^74^, rabbit antisera raised against the C-terminus of Cho1p ^34^, rabbit anti-Kar2p ^36^ and horseradish peroxidase-conjugated (Thermo Fisher Scientific) or IRDye 800CW (LI-COR) secondary antibodies.

### Miscellaneous

Preparation of yeast cell extracts, submitochondrial fractionation, phospholipid analysis, 1D BN-PAGE, and immunoblotting were performed as described previously ^17, 29^. Immunoblots using IR 800 CW secondary antibodies were imaged using an Odyssey CLx Imaging System. Immunoblots and TLC plates were quantitated by Quantity One Software by Bio-Rad Laboratories. Statistical comparisons (ns, *P* > 0.05; 1 symbol *P* ≤ 0.05; 2 symbols *P* ≤ 0.01; 3 symbols *P* ≤ 0.001; 4 symbols *P* ≤ 0.0001) were performed by student *t* test or one-way analysis of variance (ANOVA) with Holm-Sidak pairwise comparison using SigmaPlot 11 software (Systat Software, San Jose, CA); P ≤ 0.05 were deemed significant. In some cases, replicates of samples were loaded on the same SDS-PAGE gel, and thus borders of neighboring samples may be detected on the borders of some immunoblots. All graphs show the mean ± S.E.M. At least three biological replicates represent each of the experiments performed in this study, unless otherwise indicated.

## ACKNOWLEDGEMENTS

We would like to thank Drs. Carla Koehler (UCLA), Cathy Clarke (UCLA), Martin Ott (Stockholm University), Susan Michaelis (Johns Hopkins University School of Medicine), and George Carman (Rutgers University) for antibodies, Drs. Ya-Wen Lu, Ouma Onguka, Seun Ogunbona, and Matthew G. Baile for technical assistance, and Erica Avery for revision of this manuscript. This work was supported by the National Institutes of Health (R01GM111548 to S.M.C. and T32GM007445 to E.C.). The authors declare no competing financial interests.

## AUTHOR CONTRIBUTIONS

E. C. and S. M. C. co-wrote and performed experiments for this manuscript. J. M. M. performed all electron microscopy sample preparation, sectioning, and imaging.

## COMPETING INTERESTS

The authors declare no competing interests in the design and/or interpretation of this study.

## MATERIALS & CORRESPONDENCE

Correspondence and material requests should be directed to S.M.C.

## Supplementary Figures

**Supplementary Fig 1.**
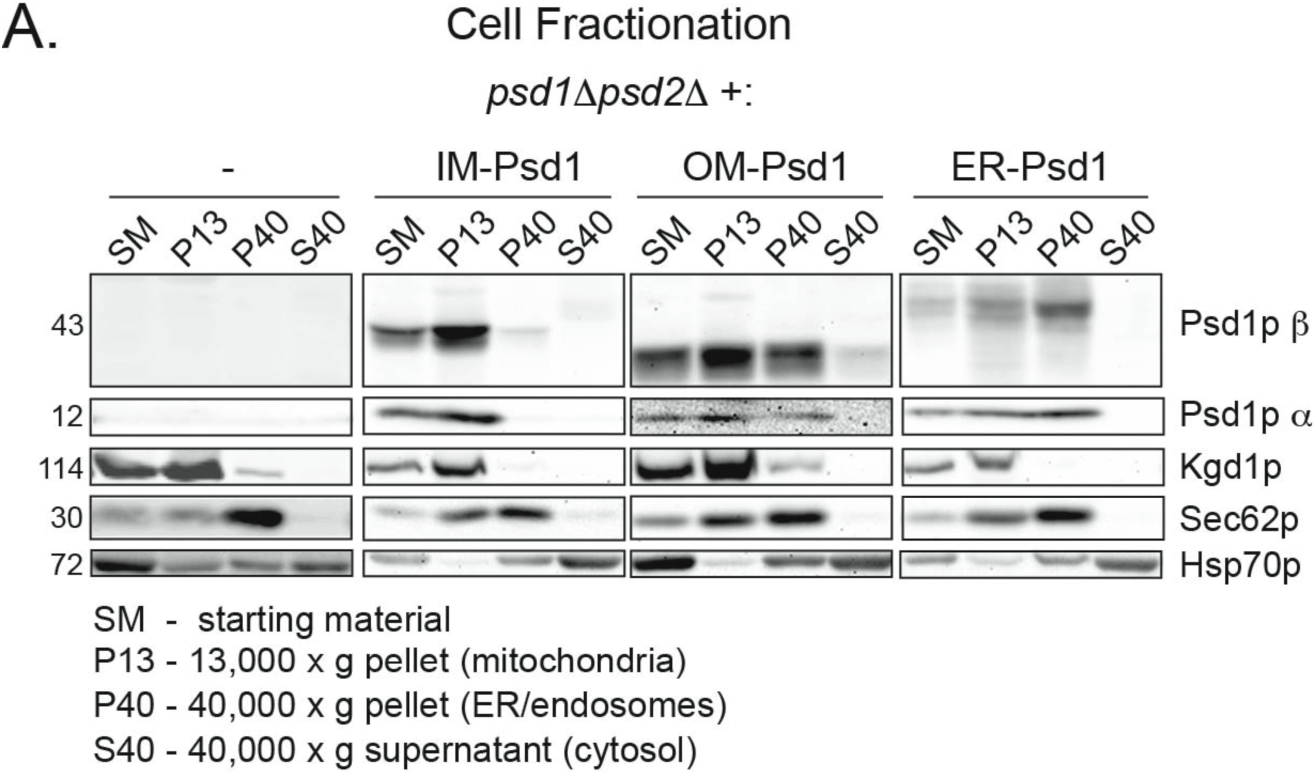
ER-Psd1p co-fractionates with ER/endosomal compartment. (A) Following growth in rich lactate medium to late log phase, fractions were collected from the indicated yeast strains by differential gravity centrifugation. Equal protein amounts from each collected fraction were resolved by SDS-PAGE and immunoblotted for Psd1p (β and α subunits) and markers for each compartment (Kgd1p for mitochondria/P13, Sec62p for the ER/P40, and Hsp70 for the cytosol/S40).

**Supplementary Fig 2.**
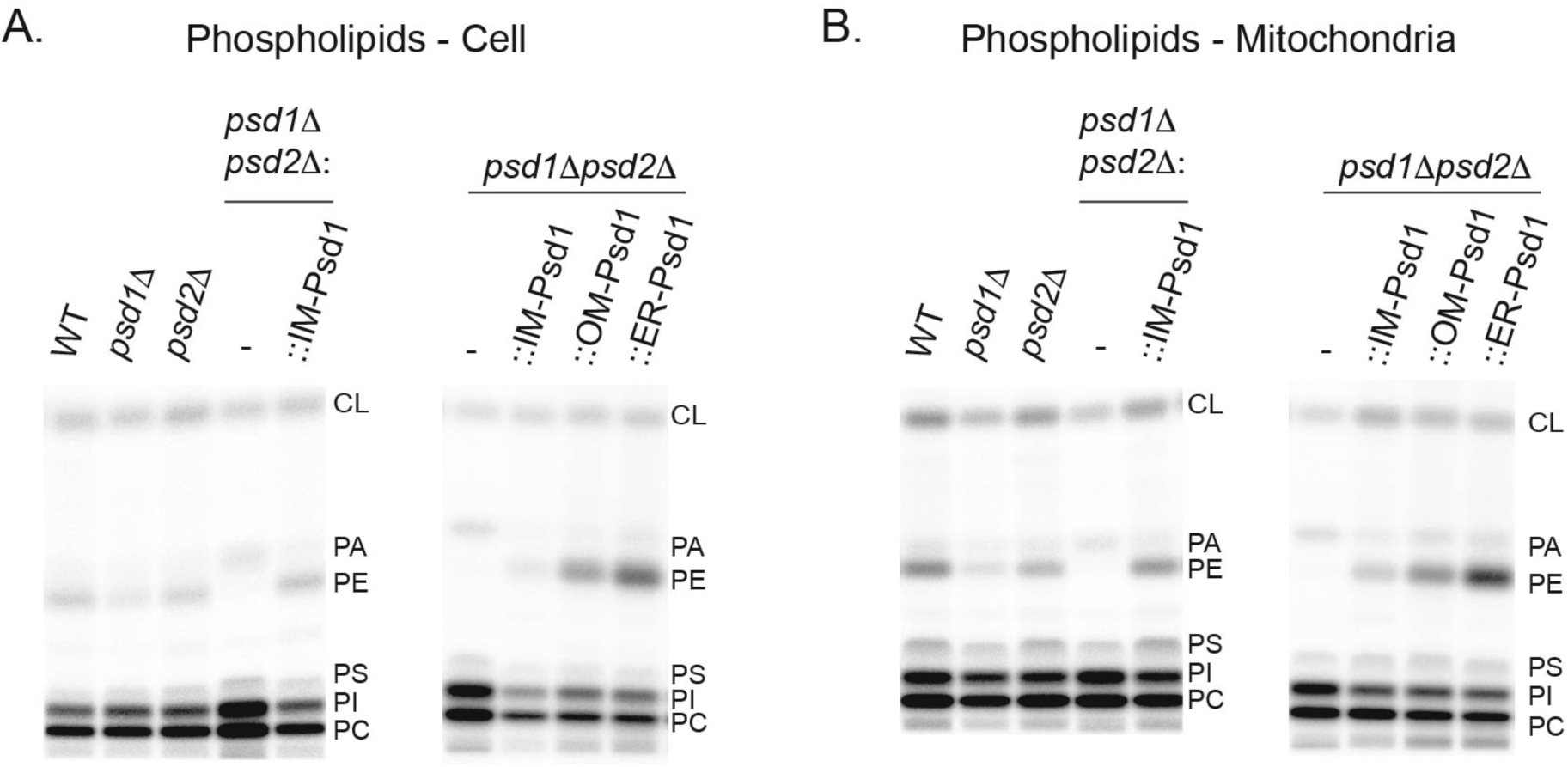
TLC images of ^32^P_i_ labeled lipids. Representative TLC plates from (A) Cellular and (B) mitochondrial lipid extractions. The migration of different phospholipid classes is indicated.

**Supplementary Fig 3.**
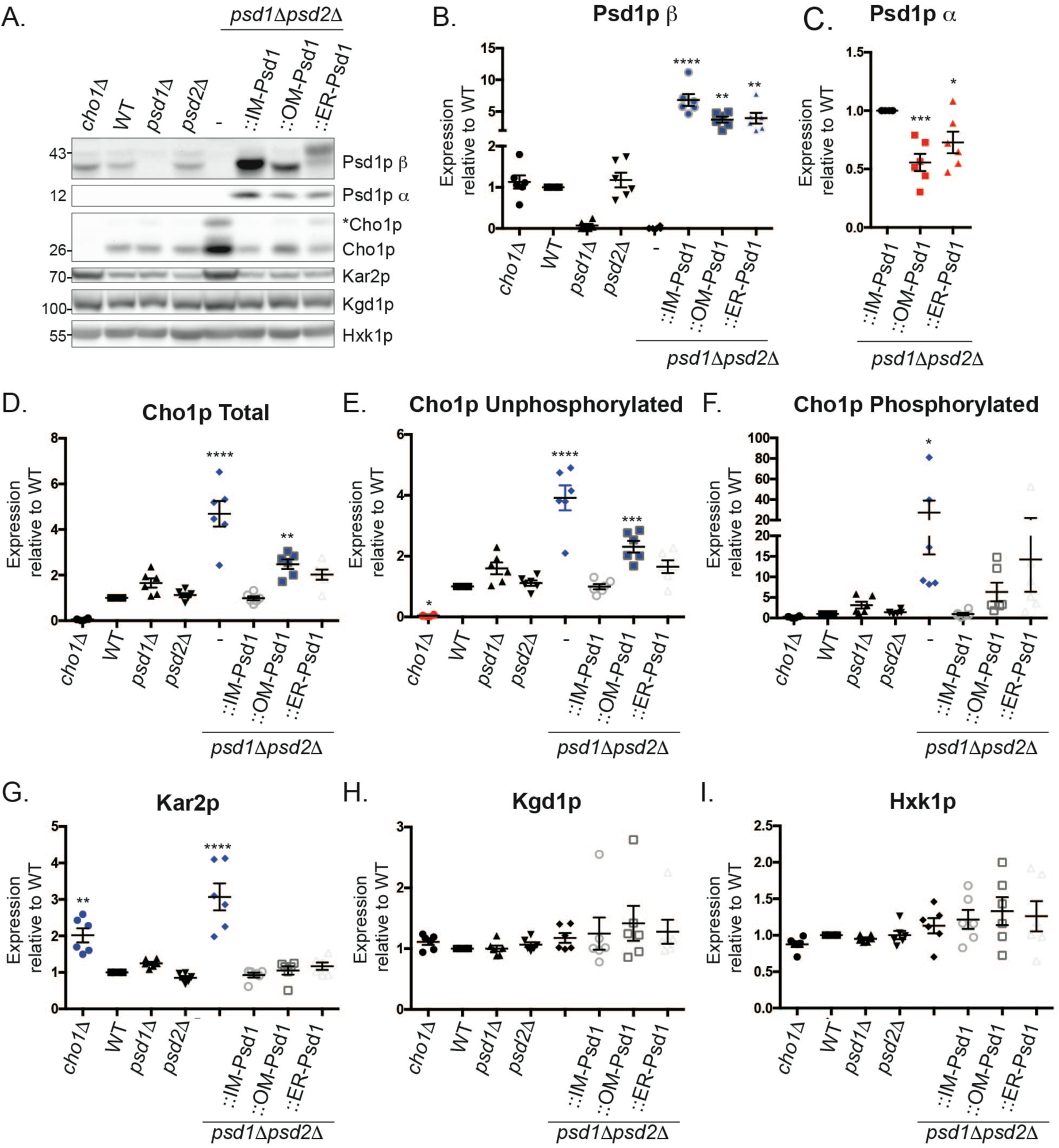
Cho1p expression and phosphorylation state is increased in *psd1*Δ*psd2*Δ yeast, which also indicate hallmarks of ER stress through elevated levels of Kar2p. Yeast strains were cultured in rich lactate medium for 2 days at 30°C. Cells were harvested by centrifugation, lysed, protein extracted, resuspended in Laemelli buffer, and analyzed by western blot. (B-I) The expression of the indicated proteins was normalized relative to WT and statistical differences compared to WT (*) determined by one-way ANOVA ± S.E.M. for n=6.

**Supplementary Fig 4.**
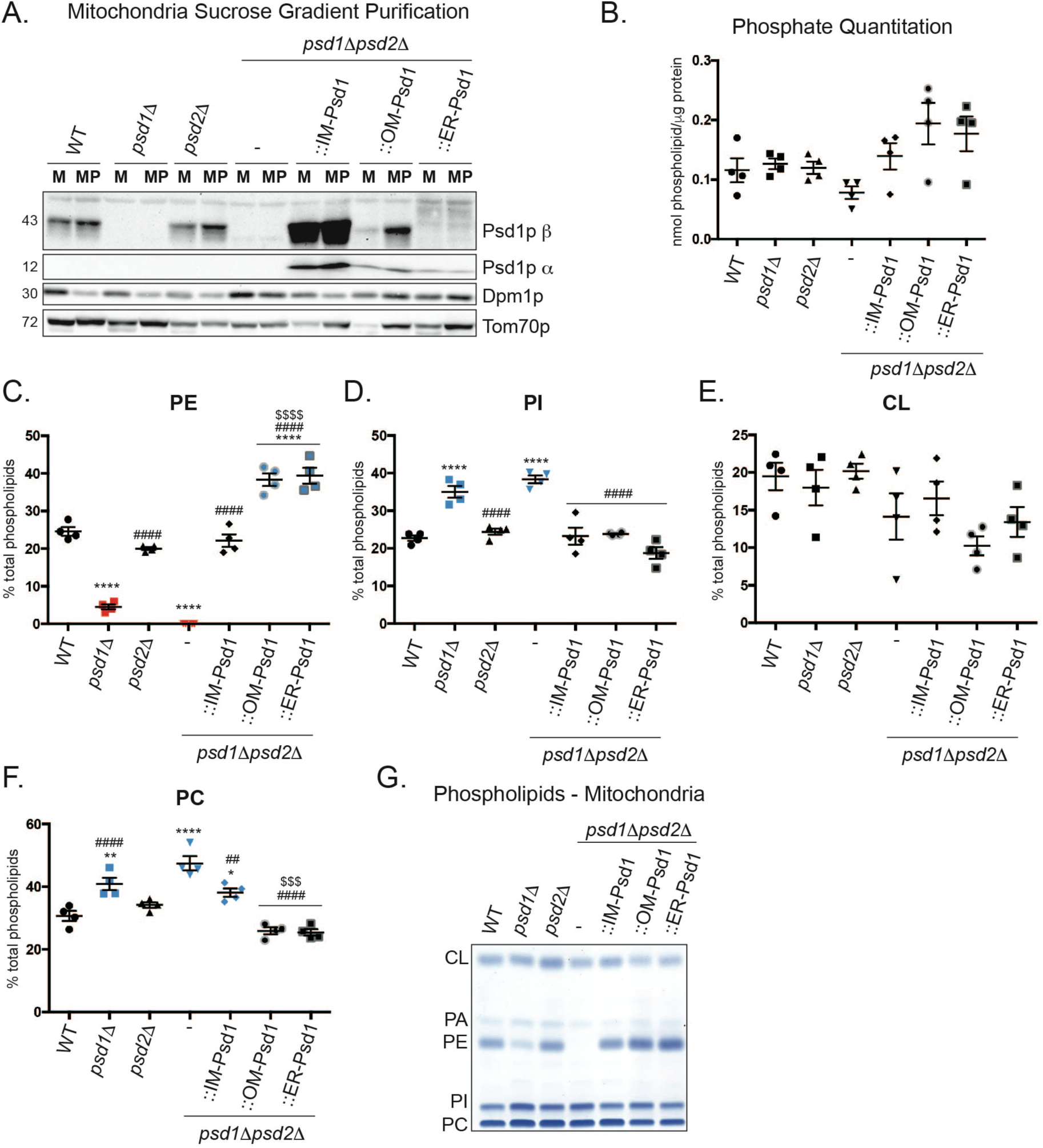
Phospholipid analysis of sucrose gradient purified mitochondria indicates equivalent levels of PE in OM-Psd1 and ER-Psd1 mitochondria. (A) After crude isolation of mitochondria by gravity centrifugation, samples were overlaid on a two-step sucrose gradient containing 32% and 60% sucrose in EM buffer and subjected to ultracentrifugation at 134,000 × *g* for 1 hour at 4°C. Purification of mitochondria was monitored by using the mitochondrial marker Tom70p and the endosomal marker Dpm1p. (B) Total phospholipid content/mitochondrial protein (mean ± S.E.M., n =4) in sucrose purified mitochondria. (C-F) Quantitation of phospholipid levels after separation by TLC and visualization by molybdenum blue staining. Analysis *versus* WT (*), *psd1*Δ*psd2*Δ (#), or IM-Psd1 ($) was performed by one-way ANOVA ± S.E.M. for n=4. (G) Representative TLC plate of phospholipid extraction from sucrose purified mitochondria followed by visualization using molybdenum blue reagent.

**Supplementary Fig 5.**
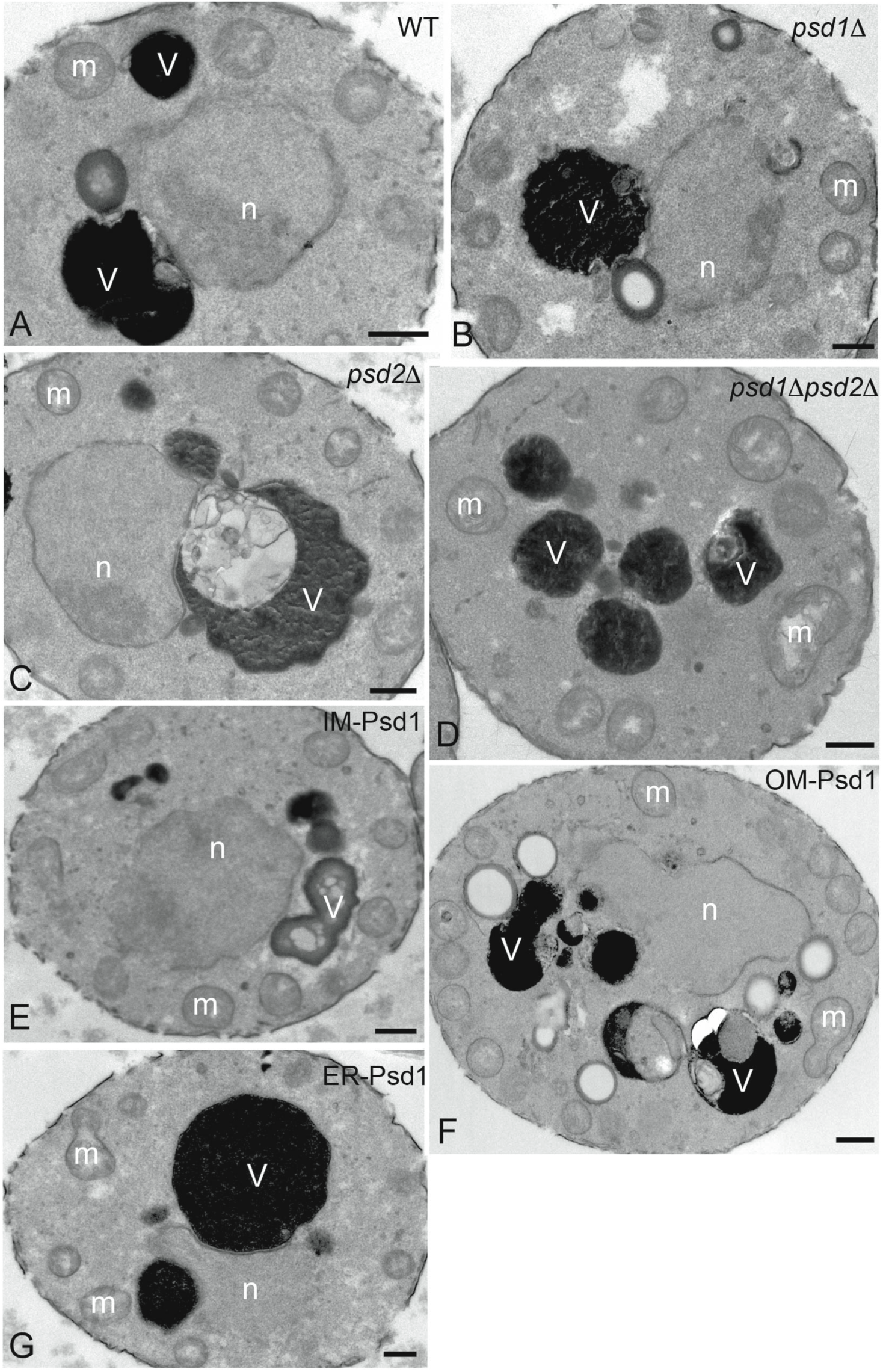
Mitochondrial morphology is not overtly affected by disruption or alteration of Psd1 pathway. Cells from the indicated strains were analyzed by transmission electron microscopy. A) GA74-1A parental wildtype strain, B) *psd1*Δ, C) *psd2*Δ, D) *psd1*Δ*psd2*Δ, E) *psd1*Δ*psd2*Δ::IM-Psd1, F) *psd1*Δ*psd2*Δ::OM-Psd1, G) *psd1*Δ*psd2*Δ::ER-Psd1. *n*, nucleus; *m*, mitochondria; and *v*, vacuole. *Bars*, 0.5 μm.

**Supplementary Fig 6.**
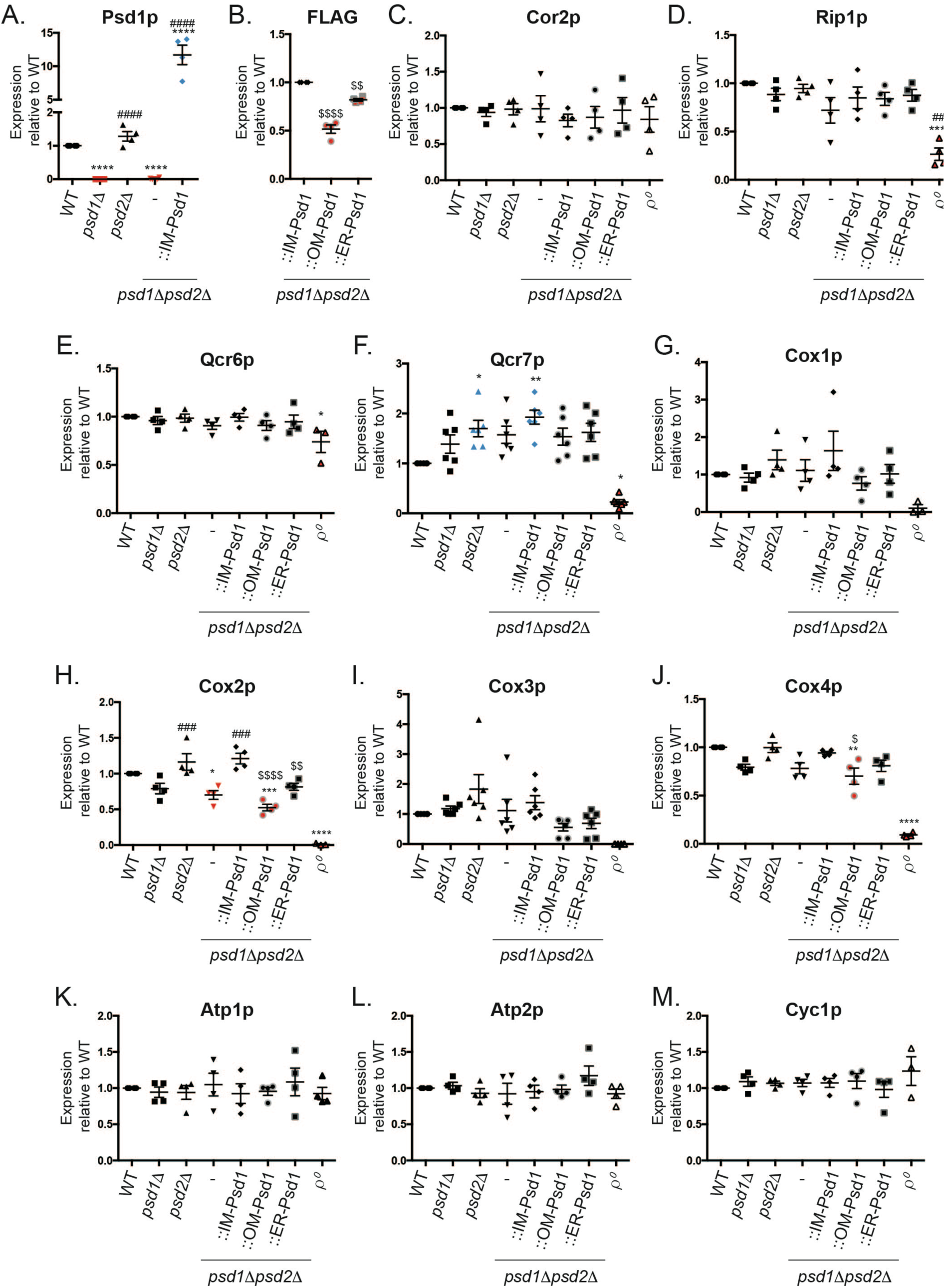
Quantitation of respiratory complex subunits. Densitometry analysis of steady state protein amounts in isolated mitochondria (30 μg) from the indicated strains (representative immunoblots shown in Fig 5C). Analysis *versus* WT (*), *psd1*Δ*psd2*Δ (#), or IM-Psd1 ($) was performed by one-way ANOVA ± S.E.M. for n=4.

**Supplementary Fig 7.**
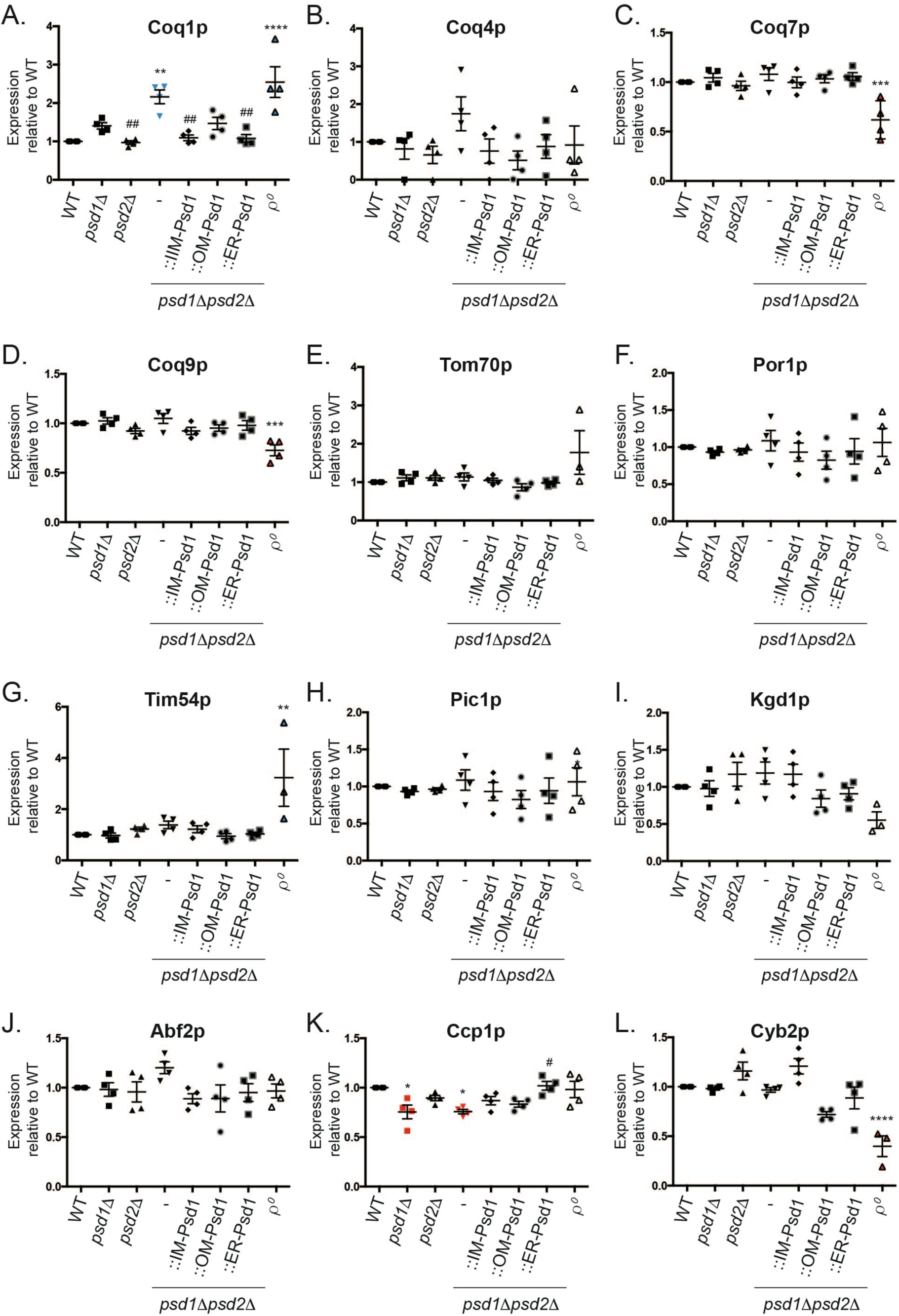
Quantitation of CoQ synthome subunits and additional mitochondrial proteins. Densitometry analysis of steady state protein amounts in isolated mitochondria (30 μg) from the indicated strains (representative immunoblots shown in Fig 5F). Analysis *versus* WT (*) or *psd1*Δ*psd2*Δ (#) was performed by one-way ANOVA ± S.E.M. for n=4.

**Supplementary Fig 8.**
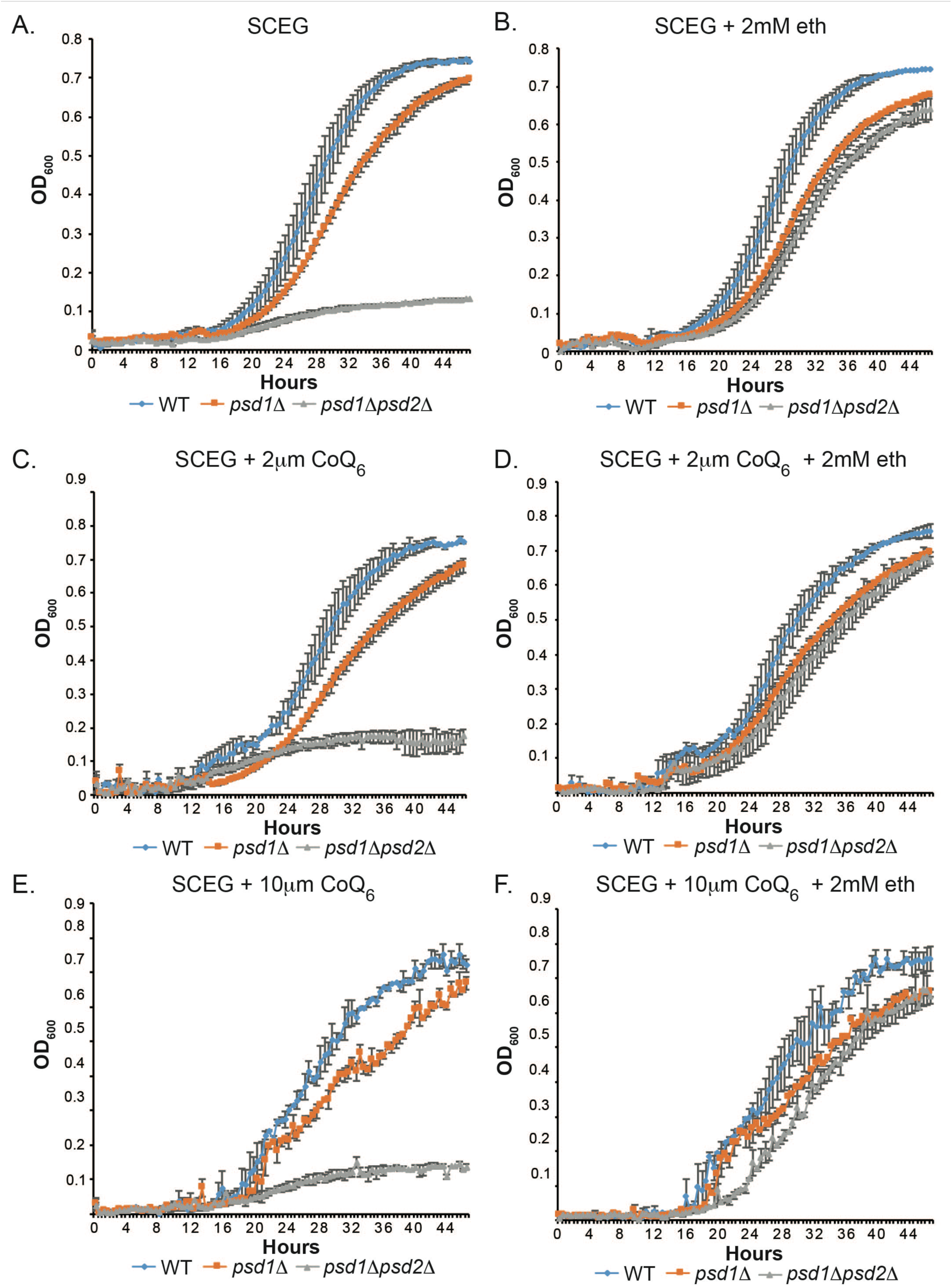
CoQ_6_ supplementation does not rescue the *psd1*Δ and *psd1*Δ *psd2*Δ growth defects on respiratory medium. OD_600_ measurements were recorded every 30 minutes for a period of 48 hours at 30°C for yeast grown in (A) SCEG, (B) SCEG + 2mM ethanolamine (eth), (C) SCEG + 2μM CoQ_6_, (D) SCEG + 2μM CoQ_6_ + 2mM ethanolamine, (E) SCEG + 10μM CoQ_6_, and (F) SCEG + 10μM CoQ_6_ + 2mM ethanolamine. ± S.E.M. for n=2.

**Supplementary Fig 9.**
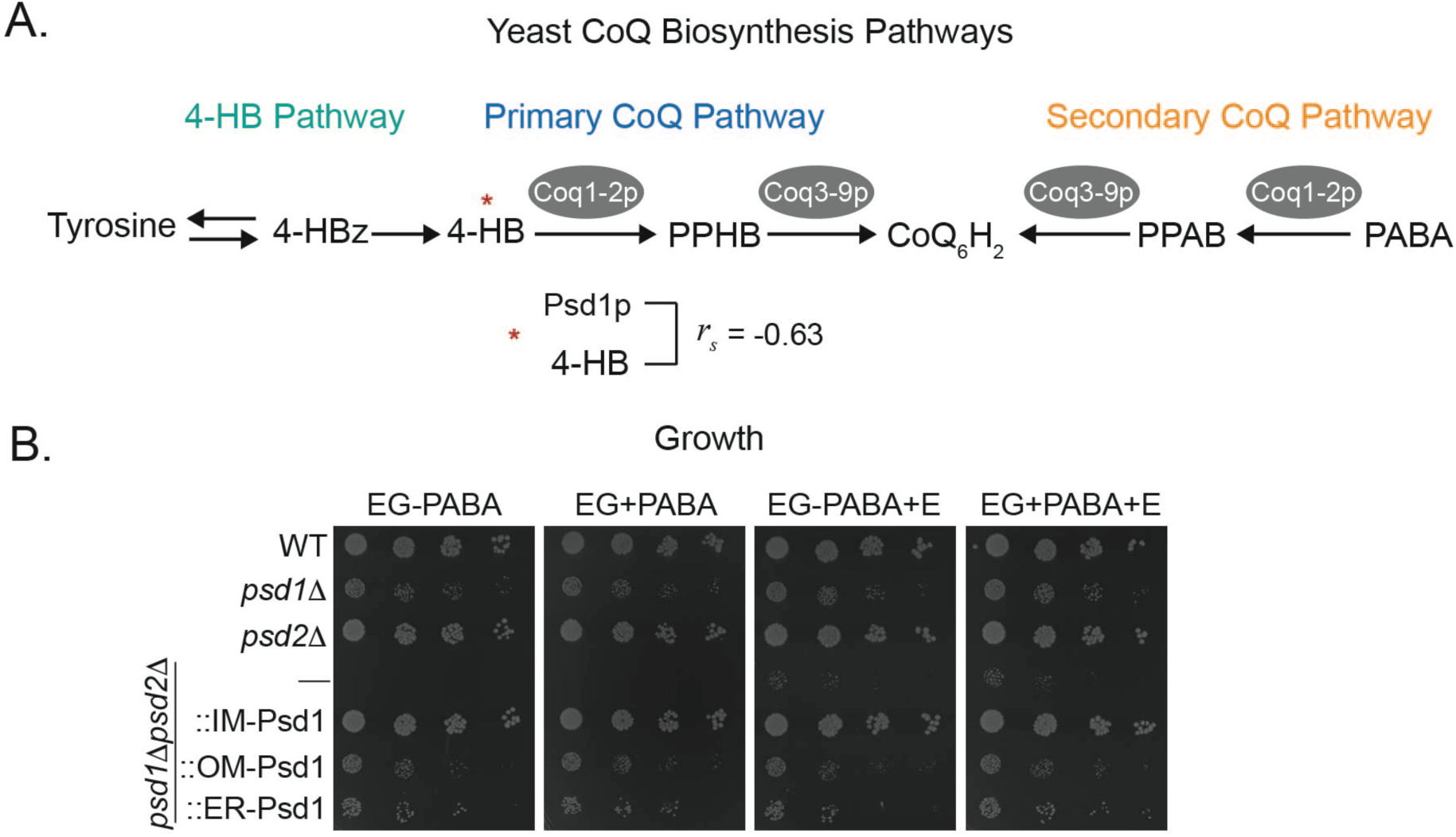
The *para*-amino benzoic acid pathway for CoQ_6_ biosynthesis is not necessary for *psd1*Δ growth. (A) Pathways for CoQ Biosynthesis; 4-hydroxy phenylpyruvate (4-HBz), 4-hydroxybenzoate (4-HB), 3-polyprenyl-4-hydroxybenzoate (PPHB), CoQ_6_H_2_, 3-hexaprenyl-4-aminobenzoate (PPAB), *para*-amino-benzoate (PABA). The indicated Spearman correlation coefficient (rs) for 4-HB molecules that share a negative correlation with Psd1p molecules, was derived from the Yeast 3 Thousand (Y3K) dataset ^41^. (B) The indicated strains were spotted and incubated at 30°C for 4 days on SCEG +/-PABA and +/- 2mM ethanolamine (+E).

**Supplementary Fig 10.**
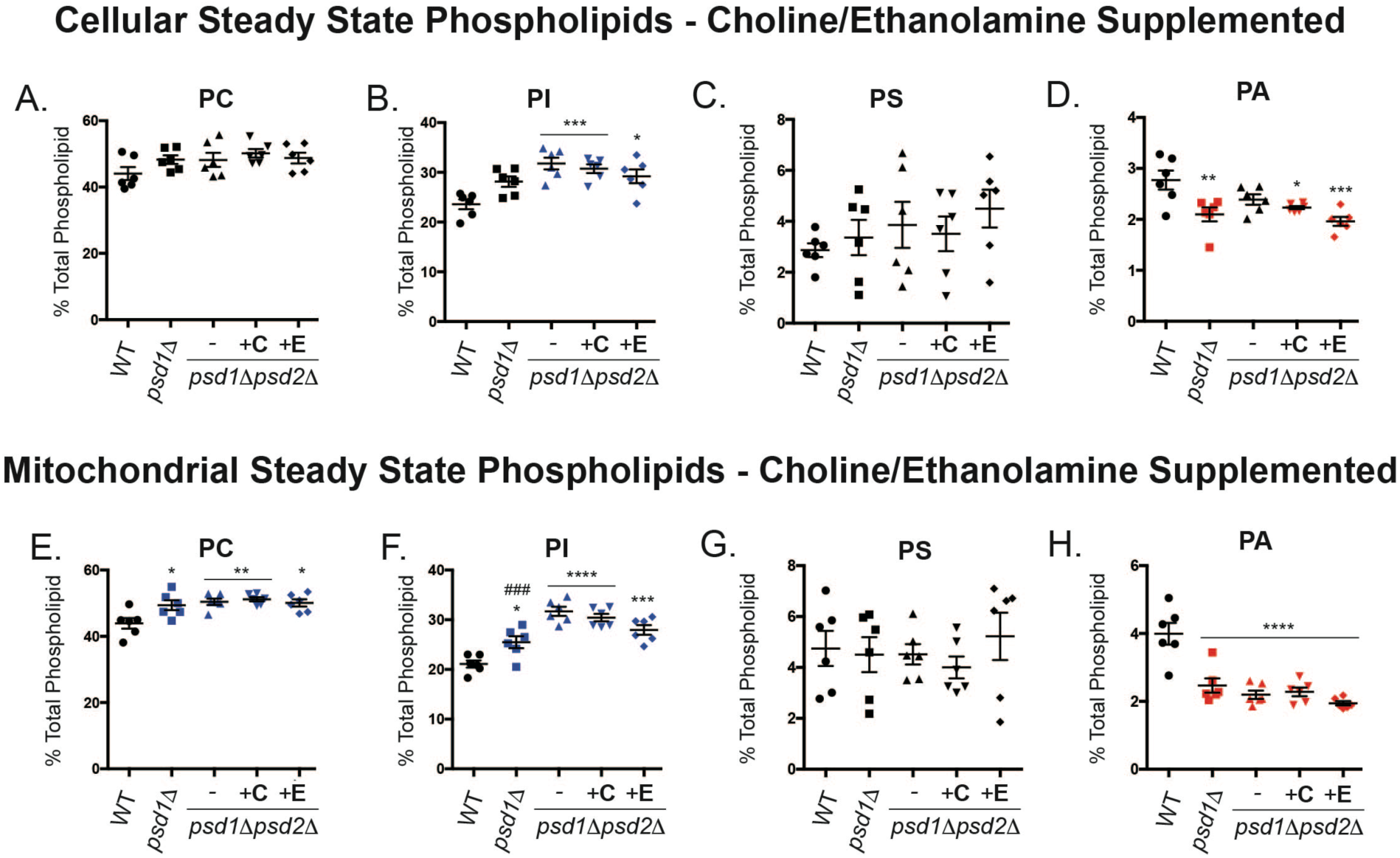
Cellular and mitochondrial phospholipid profiles from choline and ethanolamine supplemented *psd1*Δ*psd2*Δ yeast. (A-D) Cellular and (E-H) mitochondrial phospholipids from the indicated strains were labeled overnight with ^32^P_i_ and separated by TLC. All graphs show the mean ± S.E.M. for n=6 biological replicates. Significant differences *versus* WT (*) or *psd1*Δ*psd2*Δ (#) were calculated by one-way ANOVA. Key for number of symbols represented for statistical analysis interpretation 1 symbol = p < 0.05, 2 symbols = p < 0.01, 3 symbols = p < 0.001, and 4 symbols = p < 0.0001.

